# A third SARS-CoV-2 mRNA vaccine dose in people receiving hemodialysis overcomes B cell defects but elicits a skewed CD4^+^ T cell profile

**DOI:** 10.1101/2022.09.05.506622

**Authors:** Gérémy Sannier, Alexandre Nicolas, Mathieu Dubé, Lorie Marchitto, Manon Nayrac, Olivier Tastet, Alexandra Tauzin, Raphaël Lima-Barbosa, Mélanie Laporte, Rose Cloutier, Alina Sreng Flores, Marianne Boutin, Shang Yu Gong, Mehdi Benlarbi, Shilei Ding, Catherine Bourassa, Gabrielle Gendron-Lepage, Halima Medjahed, Guillaume Goyette, Nathalie Brassard, Gloria-Gabrielle Ortega-Delgado, Julia Niessl, Laurie Gokool, Chantal Morrisseau, Pascale Arlotto, Norka Rios, Cécile Tremblay, Valérie Martel-Laferrière, Alexandre Prat, Justin Bélair, William Beaubien-Souligny, Rémi Goupil, Annie-Claire Nadeau-Fredette, Caroline Lamarche, Andrés Finzi, Rita S. Suri, Daniel E. Kaufmann

**Affiliations:** Centre de Recherche of the Centre Hospitalier de l’Université de Montréal, Montreal, QC H2X 0A9, Canada; Université de Montréal, Montreal, QC H3T 1J4, Canada; Department of Microbiology and Immunology, McGill University, Montreal, QC H3A 2B4, Canada; Research Institute of the McGill University Health Centre, Montreal, QC H3H 2L9, Canada; Département de Neurosciences, Université de Montréal, Montreal, QC H3T 1J4, Canada; Nephrology Division, Centre Hospitalier de l’Université de Montréal, Montreal, QC H3X 3E4, Canada; Centre de Recherche of the Hôpital du Sacré-Cœur de Montréal, Montreal, QC H4J 1C5, Canada; Faculty of Medicine, Université de Montréal, Montréal, QC H3T 1J4, Canada; Centre de Recherche of the Hôpital Maisonneuve-Rosemont, Montréal, QC H1T 2M4, Canada; Division of Nephrology, Department of Medicine, McGill University, Montréal, QC H3G 2M1, Canada

## Abstract

Cellular immune defects associated with suboptimal responses to SARS-CoV-2 mRNA vaccination in people receiving hemodialysis (HD) are poorly understood. We longitudinally analyzed antibody, B cell, CD4^+^ and CD8^+^ T cell vaccine responses in 27 HD patients and 26 low-risk control individuals (CI). The first two doses elicit weaker B cell and CD8^+^ T cell responses in HD than in CI, while CD4^+^ T cell responses are quantitatively similar. In HD, a third dose robustly boosts B cell responses, leads to convergent CD8^+^ T cell responses and enhances comparatively more Thelper (T_H_) immunity. Unsupervised clustering of single-cell features reveals phenotypic and functional shifts over time and between cohorts. The third dose attenuates some features of T_H_ cells in HD (TNFα/IL-2 skewing), while others (CCR6, CXCR6, PD-1 and HLA-DR overexpression) persist. Therefore, a third vaccine dose is critical to achieve robust multifaceted immunity in hemodialysis patients, although some distinct T_H_ characteristics endure.

## INTRODUCTION

Implementation of SARS-CoV-2 vaccination has led to a sharp decrease in the severity of COVID-19 disease worldwide (Dagan et al., 2021; Haas et al., 2021; Tenforde et al., 2021). In the general population, mRNA vaccines against SARS-CoV-2 induce robust responses of both humoral (Earle et al., 2021; Gilbert et al., 2022; Tauzin et al., 2022a) and cellular immunity, which is dominated by B cells and Thelper (T_H_) responses with a weaker CD8^+^ T cell component (Lederer et al., 2020; Nayrac et al., 2022; Painter et al., 2021; Sahin et al., 2020). The initial series of mRNA vaccines comprised two doses. A third dose was recommended to offset waning immunity and improve recognition of variants of concern (VOCs), including Omicron (Gruell et al., 2022; Perez-Then et al., 2022; Tauzin et al., 2022b). In low-risk populations, substantial protection is conferred by one dose (Polack et al., 2020), with notable antigen-specific immunity (Baden et al., 2021; Skowronski and De Serres, 2021; Tauzin et al., 2021). Some public health agencies delayed the recommended interval between doses to increase population coverage during initial vaccine scarcity (Paltiel et al., 2021; Tuite et al., 2021). Studies in low-risk individuals subsequently showed that a longer interval between the first two doses enhanced humoral responses (Grunau et al., 2022; Hall et al., 2022; Parry et al., 2022; Tauzin et al., 2022a), and increased specific B cell responses and maturation, with little impact on T cells (Hall et al., 2022; Nicolas et al., 2022; Payne et al., 2021).

Patients with end-stage kidney disease receiving hemodialysis (HD) are susceptible to infections and demonstrate suboptimal responses to standard vaccinations against Diphteria, hepatitis B virus (HBV) or influenza (Zimmermann and Curtis, 2019). They display altered immune functions affecting B and T lymphocytes (Betjes, 2020), monocytes (Girndt et al., 2020), dendritic cells (Kim et al., 2017) and neutrophils (Kim et al., 2017) due to uremia toxins (Cohen, 2020) and blood-membrane interactions during the dialysis process (Abdelrasoul et al., 2021). However, multiple and/or higher vaccine doses proved to be an effective strategy, *e.g*. for HBV (Wu et al., 2021; Yao et al., 2021) or influenza vaccination (DiazGranados et al., 2014; Izurieta et al., 2015).

HD patients are vulnerable to SARS-CoV-2 infection, severe COVID-19 (Clark et al., 2020; Hsu et al., 2021; Williamson et al., 2020) and breakthrough events (Anand et al., 2021). Therefore, HD are considered as a high-priority population for SARS-CoV-2 vaccination. Vaccination in HD generated anti-SARS-CoV-2 antibodies, but at lower levels compared to global population (Goupil et al., 2021; Grupper et al., 2021; Simon et al., 2021), and with earlier decline (Alcazar-Arroyo et al., 2021), even after three doses (Bensouna et al., 2022; Dekervel et al., 2021; Ducloux et al., 2021). While vaccine studies in HD have focused on humoral responses, a better understanding of specific B and T cell immunity is essential to identify underlying defects. CD4^+^ T cell help plays a critical role in the generation and maintenance of adaptive immunity, particularly of B cell responses (Crotty, 2019), and CD8^+^ T cells may play a direct protective role against the virus. Some studies have shown lower SARS-CoV-2-reactive IFNγ-producing T cell frequencies in HD (Broseta et al., 2021; Melin et al., 2021; Simon et al., 2022), strengthening the need to understand this arm of the immune system. While studies suggest that long interval vaccine regimens are not appropriate for HD, resulting in weaker antibody levels, the impact on cellular immunity remains to be defined.

Herein, we conducted a prospective longitudinal cohort study to define the quantitative and qualitative trajectories of vaccine-induced antibody, B, CD4^+^ and CD8^+^ T cell responses in SARS-CoV-2 naïve HD patients receiving three mRNA SARS-CoV-2 vaccine doses, compared to antigen-specific responses in low-risk control individuals.

## RESULTS

### Study participants

We assessed cellular and antibody responses in blood samples from three cohorts of SARS-CoV-2 naïve participants who received three mRNA vaccine doses (Fig. 1A, Table 1): i) 20 people on hemodialysis (HD_S_ cohort) who received the first two doses with 5-week interval (median [interquartile range; IQ] = 35 [33-35] days); ii) 26 low-risk health care workers with no major kidney disease or immunosuppressive condition (Control Individuals, CI) who received the first two doses at a 16-week interval (median [IQ] = 111 [109-112] days), in agreement with the Quebec Public Health guidelines at the time of the study; iii) 7 HD who received a 12-week delayed second dose (median [IQ] = 83 [82-84] days) (HD_L_ long-interval HD cohort). The HD_S_ and CI cohorts were studied in detail, while we performed focused analyses on the HD_L_ cohort.

**Figure 1.**
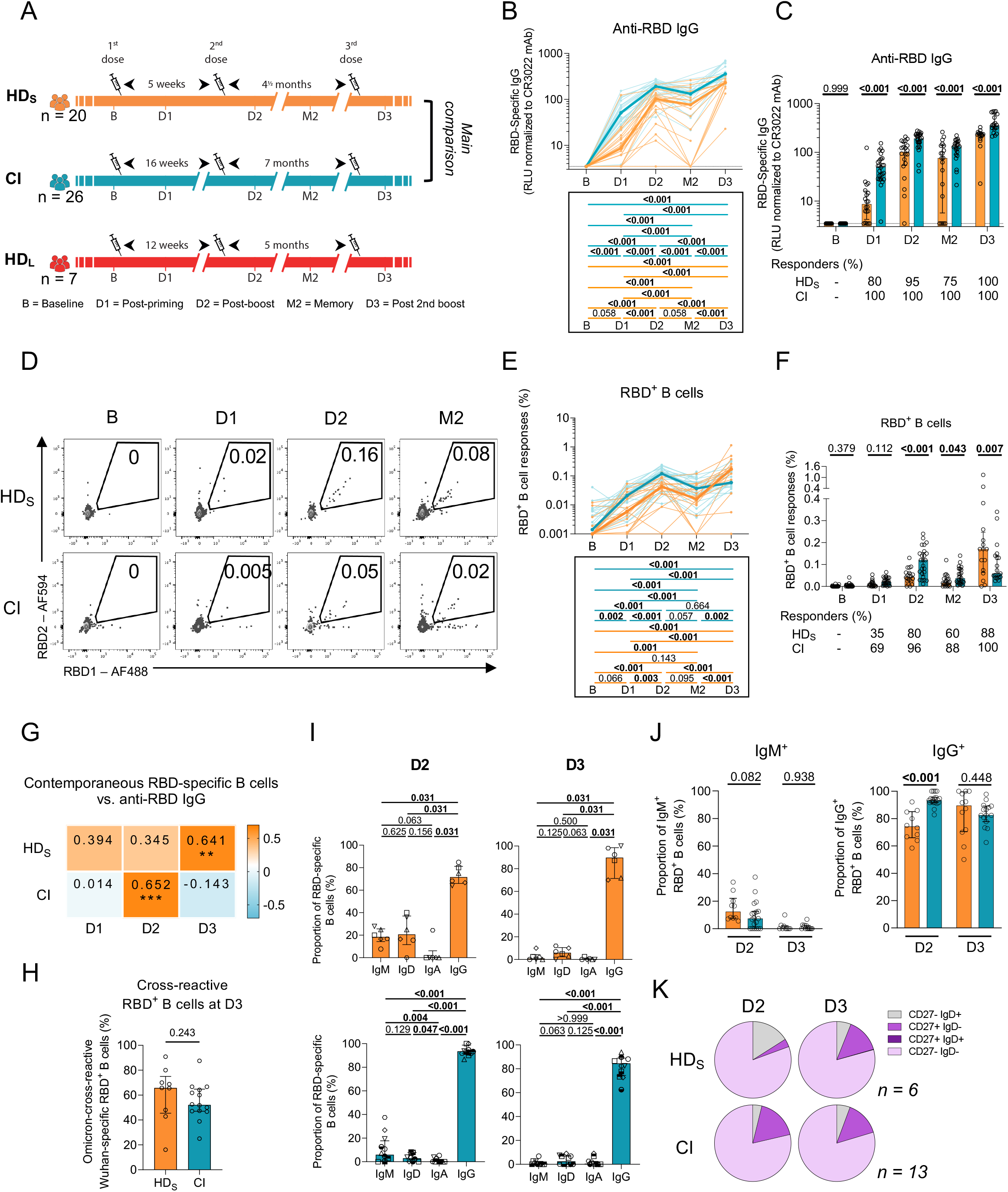
In HD patients, the initial two mRNA vaccine inoculations elicit poor B cell responses which are reinvigorated by a third dose. **(A)** Schematic representation of study design, visits, and vaccine dose administration (indicated by a syringe). Blood samples were collected at 5 time points: at baseline (B); 3-4 weeks after the first dose (D1) and the second dose (D2); 12-16 weeks after the second dose (M2); 4 weeks after the third dose (D3). Following provincial vaccination guidelines, 20 hemodialysis participants (HD_S_) received the two doses at 5 weeks interval and 26 control individuals (CI) received the two doses at 16 weeks interval. A second group of 7 hemodialysis participants (HD_L_) received a delayed second dose with an interval of 12 weeks. Actual times are summarized in Table 1. **(BC)** Kinetics of RBD^+^ IgG responses in HD_S_ participants (orange) or CI (blue) participants. **(B)** Lines connect datapoints from the same donor. The bold lines represent the median values of each cohort. Lower panel: statistical comparisons using a linear mixed model. **(C)** Comparisons between HD_S_ and CI participants. Bars represent median ± interquartile range. Intercohort statistical comparisons using a linear mixed model are shown. **(D)** Gating strategy to identify RBD^+^ B cells **(EF)** Kinetics of RBD^+^ B cell responses in HD_S_ participants (orange) or CI (blue) participants. **(E)** Lines connect datapoints from the same donor. The bold lines represent the median values of each cohort. Lower panel: statistical comparisons using a linear mixed model. **(F)** Comparisons between HD_S_ and CI participants. Bars represent median ± interquartile range. Intercohort statistical comparisons using a linear mixed model are shown. **(G)** Contemporaneous correlation of RBD^+^ B cells and anti-RBD IgG. Values and colors represent Spearman r, asterisks indicate p values (* p < 0.05, ** p < 0.01, *** p < 0.001). **(H)** Comparison between HD_S_ and CI of the proportions of Omicron-RBD^+^ B cells among wild-type Wuhan-1-specific RBD^+^ B cells. Mann-Whitney tests are shown. **(I)** Proportions of IgD, IgM, IgA, and IgG-positive cells in RBD^+^ B cells at D2 and D3 in HD_S_ and CI participants, with Wilcoxon tests. **(J)** Comparison of IgM^+^ and IgG^+^ RBD^+^ B cells between HD_S_ and CI participants at D2 and D3. Mann-Whitney tests are shown. **(K)** Proportion of IgD^+/-^ and CD27^+/-^ populations in RBD^+^ memory B cells in HD_S_ and CI participants at D2 and D3. In **BCEFG**) n=20 HD_S_, n=26 CI; in **H**) n=9 HD_S_, n=14 CI; in **IJK**) n=6 HD_S_, n=13 CI.

**Table 1.** Clinical characteristics of study participants^†^.

|  | Control Individuals (CI) | Hemodialysis Participants (HD) |  |
| --- | --- | --- | --- |
| mRNA vaccine | Long delay (16 weeks) | Short delay (HD<sub>S</sub>) (5 weeks) | Long delay (HD<sub>L</sub>) (12 weeks) |
| Variable | (n=26) | (n=20) | (n=7) |
| Vaccine regimen |  |  |  |
| Pfizer BNT162b2 vaccine (3 doses) | n=25 | n=19 | n=7 |
| Heterologous vaccine strategy (Moderna mRNA-1273 and Pfizer BNT162b2) | n=1 | n=1 | n=0 |
| Age | 51 (41-56) | 61 (55-64) | 66 (55-77) |
| Sex |  |  |  |
| Male | 11 (42%) | 13 (65%) | 4 (57%) |
| Female | 15 (58%) | 7 (35%) | 3 (43%) |
| Time on hemodialysis |  |  |  |
| Years on hemodialysis | NA | 4.9 (2.6-13.3) | 3.9 (2.8-6.1) |
| Vaccine dose spacing |  |  |  |
| Days between doses 1 and 2 | 111 (109-112) | 35 (33-35) | 83 (82-84) |
| Days between doses 1 and 3 | 329 (323-334) | 168 (166-168) | 230 (229-231) |
| Days between doses 2 and 3 | 219 (211-222) | 133 (133-133) | 147 (147-147) |
| Visits for immunological profiling |  |  |  |
| B, days before first dose | 1 (0-5) | 12 (7-12) | 1 (01-2) |
| D1, days after first dose | 21 (19-26) | 28 (28-30) | 28 (28-29) |
| D1, days before second dose | 90 (85-92) | 5 (5-7) | 54 (54-56) |
| D1, days before third dose | 306 (302-310) | 138 (138-139) | 203 (201-203) |
| D2, days after first dose | 133 (130-139) | 63 (63-63) | 111 (110-112) |
| D2, days after second dose | 21 (20-27) | 28 (28-29) | 28 (28-28) |
| D2, days before third dose | 196 (193-197) | 105 (103-105) | 119 (119-119) |
| M2, days after first dose | 224 (222-228) | 119 (117-119) | 167 (167-168) |
| M2, days after second dose | 112 (110-119) | 84 (84-84) | 84 (84-84) |
| M2, days before third dose | 104 (101-112) | 49 (49-49) | 63 (63-63) |
| D3, days after first dose | 362 (355-364) | 198 (198-198) | 265 (264-266) |
| D3, days after second dose | 249 (245-252) | 163 (163-163) | 182 (182-182) |
| D3, days after third dose | 29 (25-34) | 30 (30-32) | 35 (35-35) |

<sup>†</sup> Values displayed are medians, with IQR: interquartile range in parentheses for continuous variables, or percentages for categorical variables.
\* Control Individuals (CI), short-interval (HD<sub>S</sub>) and long-interval (HD<sub>L</sub>) hemodialysis participants cohorts were compared by: Mann-Whitney test for continuous variables, Fisher's test for categorical variables.

Blood was sampled at baseline (B) 1-12 days before the first dose; 3-4 weeks after the first dose (D1); 3-4 weeks after the second dose (D2); 3-4 months after the second dose (memory; M2) and 4 weeks after the third dose (D3). Donors with breakthrough COVID-19 events were excluded afterwards. There were no significant differences between HD_S_ and HD_L_ in term of sex, age, nor time on hemodialysis. HD_S_ and HD_L_ were respectively 10 and 15 years-older than the CI. Time prior B, between D1 and D2, and between injection and sampling were significantly different.

### In HD patients, the initial two mRNA vaccine inoculations elicit poor B cell responses which are reinvigorated by a third dose

We measured the levels of IgG targeting the receptor binding domain (RBD). Some antibodies against RBD are neutralizing (Ju et al., 2020; Piccoli et al., 2020; Shi et al., 2020; Wu et al., 2020) and associated with vaccine efficacy (Barin et al., 2022). At baseline anti-RBD IgG were undetectable in all participants, consistent with their SARS-CoV-2 naïve status (Fig. 1BC). Both cohorts developed anti-RBD IgG responses after each vaccine dose (D1, D2 and D3) with a small decline at a memory timepoint (M2) (Fig. 1B). However, the antibody levels at D1 were lower in HD_S_ compared to CI, with a median-fold difference of 6. The antibody levels in HD_S_ remained significantly lower through all follow-up timepoints (Fig. 1C). Only 75% (15/20) of HD_S_ seroconverted after the first dose compared to 96% (25/26) of CI. However, all HD_S_ experienced an increase in anti-RBD IgG responses after the third dose (Fig. 1C). In the HD_L_ regimen, the level of anti-RBD IgG were not significantly different compared to HD_S_ (Fig. S1A), suggesting that unlike low-risk populations (Chatterjee et al., 2022; Hall et al., 2022; Parry et al., 2022; Tauzin et al., 2022a), longer interval regimens are not beneficial in HD.

We next measured RBD^+^ B cells using two fluorescently labeled recombinant RBD probes (Fig. 1D, Fig. S1B) (Nayrac et al., 2022; Nicolas et al., 2022). We observed differences in both magnitude and longitudinal trajectories of B cell responses between cohorts (Fig. 1EF). There was a trend for weaker priming of B cell responses (D1) in HD_S_ than CI which did not reach statistical significance after correction for multiple comparisons. At D2 and M2, the frequencies of RBD^+^ B cells in HD_S_ consistently trailed those in CI, resulting in almost parallel curves (Fig. 1E). In contrast, the responses to the third dose differed, with more robust expansion of B cell responses in HD_S_ compared CI. Consequently, we observed a stronger B cell responses at D3 in HD_S_ than in CI (Fig. 1F). Consistent with the antibodies (Fig. S1A) and unlike CI (Nicolas et al., 2022), a long interval in HD_L_ did not improve the generation of the RBD^+^ B cell pool (Fig. S1C). The delayed kinetics of anti-RBD IgG responses in HD_S_ compared to CI is illustrated by contemporaneous associations between B cell and antibody responses: while we observed in CI a significant positive correlation between RBD^+^ B cells and anti-RBD IgG at D2, this correlation only appeared at D3 in HD_S_ (Fig. 1G).

The rapid worldwide spread of the Omicron variant has decreased vaccine efficacy against infection (Bian et al., 2022; Hacisuleyman et al., 2021). However, protection against severe disease remains good, and is significantly increased by a third vaccine dose (Planas et al., 2022; Tauzin et al., 2022b; Tregoning et al., 2021). To determine if hemodialysis treatment was associated with altered viral cross-recognition by B cell, we tested if HD_S_ immunized with (WT) Wuhan-1 strain Spike could elicit cross-reactive B cell responses against Omicron RBD (Fig. S1D). Among all the WT RBD^+^ B cells at D3, 65% co-stained for Omicron RBD probes, indicating cross-reactivity (Fig. 1H). No significant difference was observed between HD_S_ and CI (Fig. 1H).

We next assessed differentiation of RBD^+^ B cells following vaccination. To avoid phenotyping bias, we only included donors in whom we detected ≥ 5 RBD^+^ B cells at every timepoint. As the rare RBD^+^ B cells in HD_S_ at D1 precluded reliable phenotyping, we focused on D2 and D3. We measured expression of IgM, IgD, IgA, and IgG on RBD^+^ B cells (Fig. S1E-G). While IgG^+^ cells were dominant in both cohorts at all timepoints (Fig. 1IJ, S1F), its fraction was lower in HD_S_ at D2, and those of IgM^+^ and IgD^+^ cells were higher (Fig 1J, Fig. S1G). The profiles converged between cohorts at D3. We next determined the memory differentiation profile of RBD^+^ B cells using IgD and CD27 co-expression (Fig. S1H). CD27 is expressed on memory B cells (Macallan et al., 2005; Tangye et al., 2003) and IgD is mostly found on unswitched B cells (Kaminski et al., 2012). In both cohorts, RBD^+^ B cells were mainly IgD^-^CD27^-^ (Fig.1K). In HD_S_, CD27^-^IgD^+^ cells represented 15% of RBD^+^ B cells at D2 and disappeared at D3 in favor of mature CD27^+^IgD^-^ cells. In CI, no such immature population was observed at D2, and frequencies of mature RBD^+^ B cells were similar at both time points. Quantitatively, memory B cells increased after the third dose in both groups (Fig. S1I).

Our data show that compared to a CI cohort, HD_S_ elicit low RBD^+^ and mature B cell responses after two doses, consistent with lower antibody levels. A third immunization in HD_S_ is critical to achieve B cell responses of higher magnitudes than those observed in CI and leads to convergent differentiation profiles.

### Vaccination induces strong CD4^+^ T cell responses but poor CD8^+^ T cell immunity in hemodialysis participants

SARS-CoV-2-specific CD4^+^ T cells help is crucial to the development of B cells responses and correlates with long-term humoral responses and CD8^+^ T cells immunity (Apostolidis et al., 2021; Goel et al., 2021; Nayrac et al., 2022; Painter et al., 2021; Sette and Crotty, 2021). We measured Spike (S)-specific T cell (Fig. S2A) responses using activation induced marker (AIM) (Morou et al., 2019; Nayrac et al., 2022; Niessl et al., 2020; Tauzin et al., 2022a) and intracellular cytokine staining (ICS) assays (Nayrac et al., 2022).

S-specific CD4^+^ and CD8^+^ T cell responses were assessed by an OR Boolean combination gating strategy of the upregulation of CD69, CD40L, 4-1BB, and OX-40 upon a 15h-stimulation with an overlapping peptide pool spanning the Spike coding sequence (Fig. S2B). This strategy detected S-specific AIM^+^ CD4^+^ (Fig. S2C) and CD8^+^ T cells responses (Fig. S2D). To assess the functionality of the specific T cells, we measured the expression of IFNγ, IL-2, TNFα, IL-17A, IL-10, and CD107a following a 6h-stimulation with the Spike peptide pool. We determined total ICS^+^ CD4^+^ T cell responses by a similar OR Boolean combination gating strategy (Fig. S2E). Specific ICS^+^ CD4^+^ T cell responses were detected at all timepoints (Fig. S2F). In most participants, no significant CD8^+^ T cell functions were detected and could not be further assessed (Fig. S2G).

AIM^+^ CD4^+^ T cell responses in HD_S_ significantly increased after priming, plateaued at D2, waned slightly at M2, and further increased to peak at D3 (Fig. 2A). In CI, the increase at D3 was more muted (Fig. 2A). We observed a trend for stronger AIM^+^ CD4^+^ T cell responses in HD_S_ than in CI at M2 and significantly higher magnitudes at D3 (Fig. 2B). ICS^+^ CD4^+^ T cell responses in HD_S_ followed trajectories comparable to the AIM responses (Fig. 2CD), with stronger effector CD4^+^ T cell responses in HD_S_ compared to CI at D3 (Fig. 2D). Unlike CD4^+^ T cells, AIM^+^ CD8^+^ T cell responses were lower in HD_S_ than CI at all timepoints except M2 and D3, at which they converged (Fig. 2EF). A trend for an increased CD8^+^ T cell responses in HD_S_ at D3 in comparison to baseline was observed (Fig 2 EF). Of note, the CI cohort was characterized by sizeable pre-existing CD8^+^ T cell responses at baseline, which likely impacted the patterns observed (Fig 2 EF). As observed with the RBD^+^ B cell responses, the long interval in HD_L_ seemed detrimental for mRNA-vaccine elicited cellular responses, neither for CD4^+^ (Fig. S2HI) nor CD8^+^ (Fig. S2J) T cell responses.

**Figure 2.**
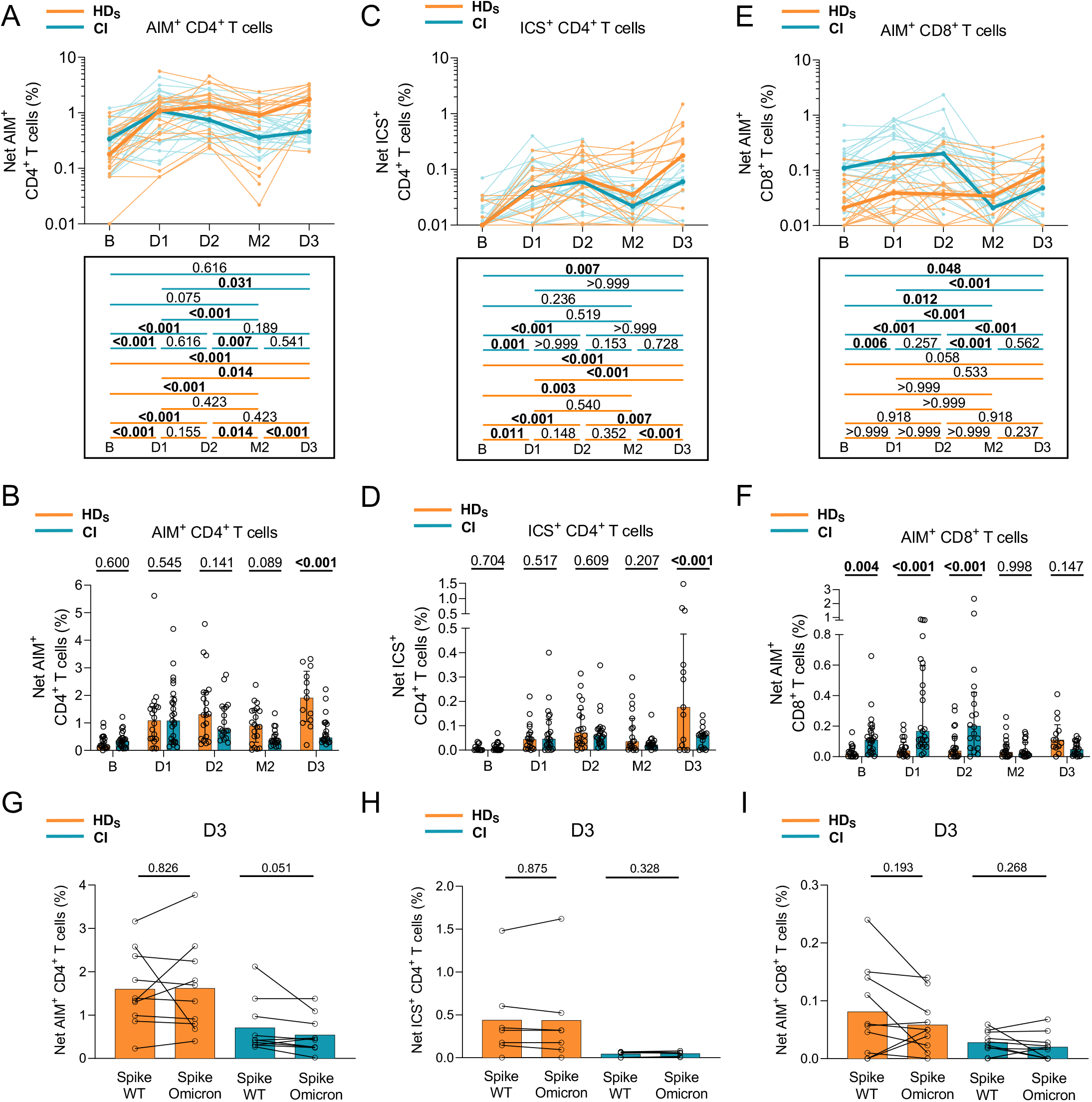
Vaccination induces strong CD4^+^ T cell responses but poor CD8^+^ T cell immunity in hemodialysis participants. Frequencies of SARS-CoV-2 S-specific CD4^+^ and CD8^+^ T cells in HD_S_ (orange) and CI (blue) who received three vaccine doses. PBMCs were stimulated *ex vivo* with a pool of overlapping Spike peptides. **(AB)** Net AIM^+^ CD4^+^ T cell responses. **(A)** Longitudinal analysis of S-specific AIM^+^ CD4^+^ T cell responses. Lines connect datapoints from the same donor. The bold lines represent the median values of each cohort. Lower panel: statistical comparisons using a linear mixed model. **(B)** Comparisons between HD_S_ and CI participants. Bars represent median ± interquartile range. Intercohort statistical comparisons using a linear mixed model are shown. **(CD)** Net ICS^+^ CD4^+^ T cell responses measured by ICS. **(C)** Longitudinal analysis of the magnitude of ICS^+^ CD4^+^ T cell responses. Lines connect datapoints from the same donor. The bold lines represent the median values of each cohort. Lower panel: statistical comparisons using a linear mixed model. **(D)** Comparisons between HD_S_ and CI participants. Bars represent median ± interquartile range. Intercohort statistical comparisons using a linear mixed model are shown. **(EF)** Net AIM^+^CD8^+^ T cell responses. **(E)** Longitudinal analysis of S-specific AIM^+^CD8^+^ T cell responses. Lines connect datapoints from the same donor. The bold lines represent the median values of each cohort. Lower panel: statistical comparisons using a linear mixed model. **(F)** Comparisons between HD_S_ and CI participants. Bars represent median ± interquartile range. Intercohort statistical comparisons using a linear mixed model are shown. **(G-I)** Comparison of WT Wuhan-1-specific and Omicron-specific CD4^+^ and CD8^+^ T cell responses in HD_S_ (orange) and CI (blue) participants. Mann-Whitney tests are shown. **(G)** Net S-specific AIM^+^ CD4^+^ T cell responses **(H)** Net S-specific ICS^+^ CD4^+^ T cell responses. **(I)** Net S-specific AIM^+^ CD8^+^ T cell responses. In **A-F**, **J**) n=20 HD_S_, n=26 CI participants. In **G-I**) n=10 HD_S_, n=10 CI participants.

We next assessed the presence of Omicron-reactive CD4^+^ and CD8^+^ T cell responses using an overlapping Omicron Spike peptide pool on D3 samples. We detected Omicron-reactive AIM^+^ CD4^+^ (Fig. 2G), ICS^+^ CD4^+^ (Fig. 2H), and AIM^+^ CD8^+^ (Fig. 2I) T cell responses in both cohorts. The magnitude of WT and Omicron S-specific CD4^+^ T cell responses did not differ between groups, suggesting cross-reactivity.

These data demonstrate the emergence of robust SARS-CoV-2-specific CD4^+^ T cell responses after the initial priming dose in HD_S_, while a minimum of three doses was required to generate low CD8^+^ T cell responses.

### mRNA vaccines elicit a multifaceted AIM^+^ CD4^+^ T cell responses with qualitative features in HD distinct from CI participants

To qualitatively evaluate the S-specific AIM^+^ CD4^+^ T cell responses we applied unsupervised analyses, as described (Nayrac et al., 2022). We studied chemokine receptors that are preferentially expressed by some lineages and involved in tissue homing (CXCR5 for T_FH_; CXCR3 for T_H1_; CCR6 for T_H17_/T_H22_ and mucosal homing; CXCR6 for pulmonary mucosal homing (Day et al., 2009; Morgan et al., 2015), activation markers (HLA-DR and CD38), and an inhibitory checkpoint (PD-1).

The distribution of clustered populations was represented by the uniform manifold approximation and projection (UMAP) algorithm (Becht et al., 2018). Cluster identity was performed using Phenograph (Levine et al., 2015), resulting in the identification of 14 clusters (Fig. 3AB) based on distinct profiles of relative marker expression (Fig. 3C and S3A). The 14 clusters were detectable at all timepoints (Fig. 3AD), most of them following frequency trajectories consistent with those observed for total AIM^+^ CD4 T cells in both cohorts (Fig. 3E, Fig. S3B). Some qualitative differences in relative proportions persisted after the third dose, with in HD_S_ significant expansion of C4 and C7, two clusters enriched in CXCR6, CCR6, CD38 and PD-1 expression. Additional upregulation of HLA-DR characterized C7 (Fig. 3F, Fig. S3C). In univariate analyses, we observed at D3 in HD_S_ significant expansion of PD-1^+^, HLA-DR^+^, CXCR6^+^ cells and, to a lesser extent, CD38^+^ cells both in absolute frequencies and as relative fractions of AIM^+^ CD4^+^ T cells (Fig. 3G, Fig. S3D).

**Figure 3.**
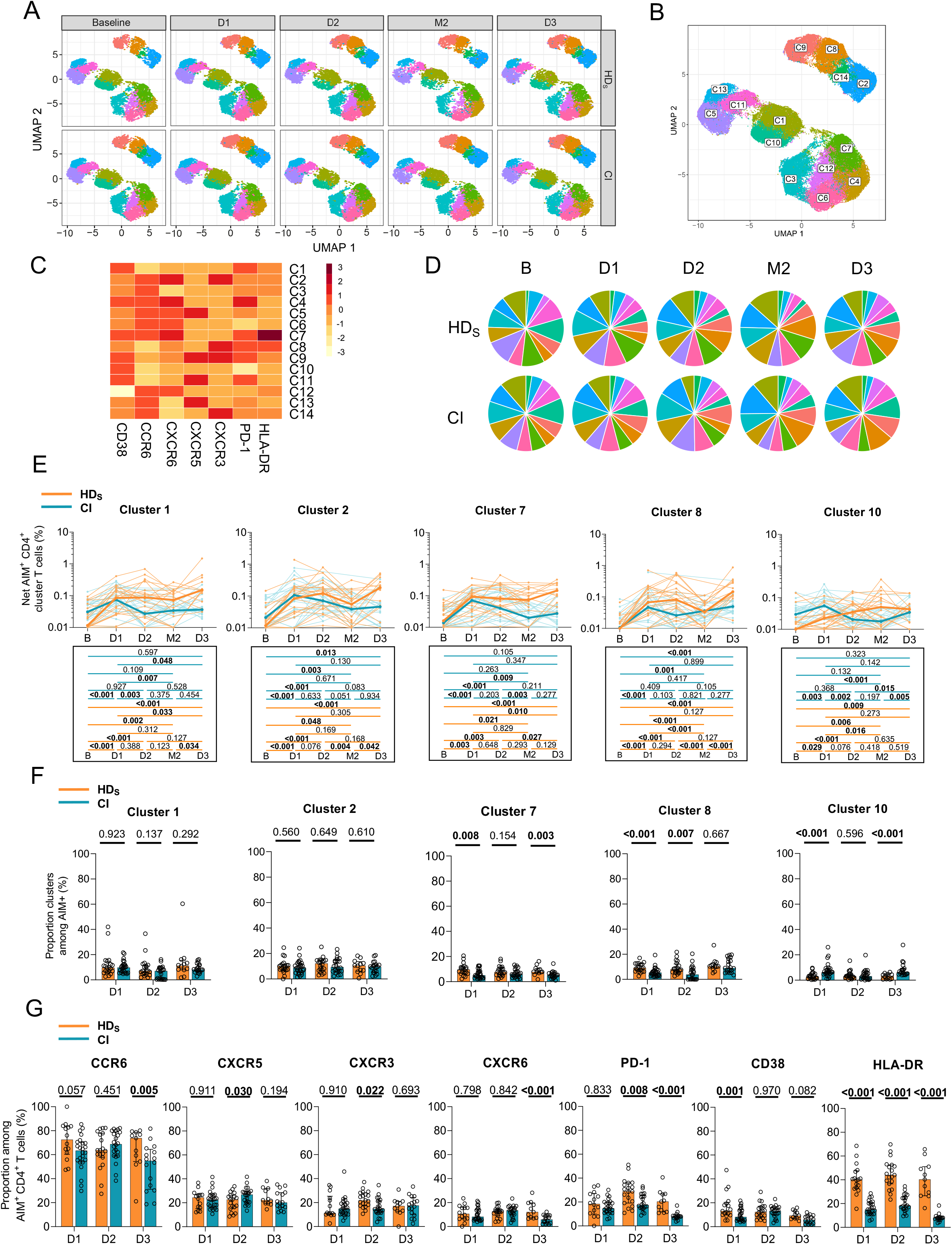
mRNA vaccines elicit a multifaceted AIM^+^ CD4^+^ T cell response with qualitative features in HD distinct from CI participants. **(A)** Multiparametric UMAP representation of S-specific AIM^+^ CD4^+^ T cells based on CD38, HLA-DR, CCR6, CXCR6, CXCR5, CXCR3, and PD-1 expression at each time point, aggregated data for the HD_S_ and CI cohorts. The colors identify 14 populations clustered by unsupervised analysis using Phenograph. **(B)** Each cluster is labeled on the global UMAP. **(C)** Heat map summarizing for each cluster the MFI of each loaded parameter. **(D)** Pie charts depicting the representation of each identified cluster within total AIM^+^ CD4^+^ T cells. **(E)** Longitudinal net frequencies of selected AIM^+^ CD4^+^ T cell clusters in HD_S_ (orange) and CI (blue) participants (left) for cluster 1, 2, 7, 8, and 10. Lines connect data from the same donor. The bold lines represent the median values of each cohort. Wilcoxon test for each pairwise comparison. **(F)** Proportions of AIM^+^ clusters 1, 2, 7, 8 and 10 among AIM^+^ CD4^+^ T cells in HD_S_ and CI at D1, D2 and D3. Bars represent median ± interquartile range. Mann-Whitey tests are shown. **(G)** Cohort comparison of univariate analyses. Bars represent median ± interquartile range. Statistical comparisons using a linear mixed model are shown. In **A-G**) n=20 HD_S_, n=26 CI participants.

Therefore, mRNA vaccination elicits in both HD_S_ and CI a multifaceted response already observed after the first dose. After the full course of three vaccinations, T_H_ responses show some qualitative differences between cohorts, with higher expression in HD_S_ of chemokine receptors associated with mucosal immunity (CCR6, CXCR6), immune activation (HLA-DR) and the inhibitory immune checkpoint PD-1.

### The first two vaccine inoculations elicit in HD_S_ a TNFα/IL-2 skewed T_H_ profile that is attenuated by the third dose

Given the qualitative differences observed with AIM assays, we used the same unsupervised approach to identify differences in CD4^+^ T cell effector functions. Expression of TNFα, CD107A, IL-10, IFNγ, IL-2 and IL-17A defined 8 functional clusters, also detected at all timepoints (Fig. 4A-C, Fig. S4A) in both cohorts (Fig. 4D). All clusters increased in magnitude after the first two doses, irrespectively of clinical status (Fig. 4E, Fig. S4B). Several qualitative differences were observed at D1 and D2 between cohorts, with the TNFα/IL-2-expressing C6 enriched in HD_S_ and C1, C2, C8 overrepresented in CI (Fig 4F). The third dose led to partial convergence of the functional profiles, although the differences in C6 and C8 proportions remained significant at D3. In univariate analyses, TNFα and IL-2 expression was also higher, and IFNγ and IL-10 lower in HD_S_ compared to CI during the initial vaccination series. Statistically significant differences were mostly abrogated by the third dose (Fig 4G, Fig. S4D).

**Figure 4.**
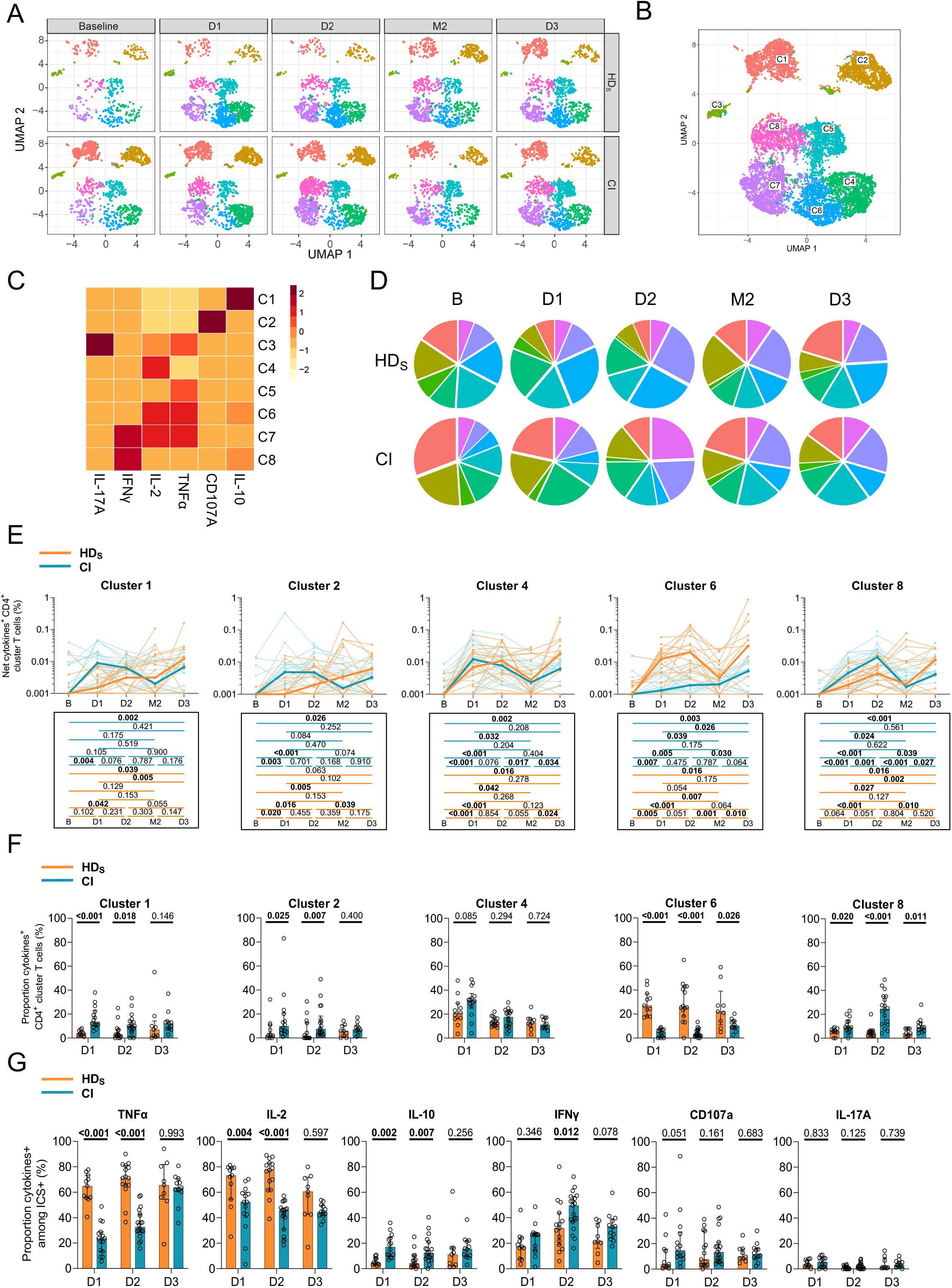
The first two vaccine inoculations elicit in HD_S_ a TNFα/IL-2 skewed T_H_ profile that is attenuated by the third dose. **(A)** Multiparametric UMAP representation of S-specific ICS ^+^ CD4^+^ T cells based on TNFα, CD107a, IL-10, IFNγ, IL-2 and IL-17A expression at each time point, aggregated data for the HD_S_ and CI cohorts. The colors identify 8 populations clustered by unsupervised analysis using Phenograph. **(B)** Each cluster is labeled on the global UMAP. **(C)** Heat map summarizing for each cluster the MFI of each loaded parameter. **(D)** Pie charts depicting the representation of each identified cluster within total ICS^+^ CD4^+^ T cells. **(E)** Longitudinal frequencies of selected ICS^+^ CD4^+^ T cell clusters 1, 2, 4, 6 and 8 in HD_S_ (orange) and CI (blue) participants. Lines connect data from the same donor. The bold lines represent the median values of each cohort. Wilcoxon tests for each pairwise comparisons are shown below the graphs. **(F)** Proportions of ICS^+^ clusters 1, 2, 4, 6 and 8 among ICS^+^ CD4^+^ T cells in HD_S_ and CI at D1, D2 and D3. Bars represent median ± interquartile range. Mann-Whitey tests are shown. **(G)** Cohort comparison of univariate analyses. Bars represent median ± interquartile range. Statistical comparisons using a linear mixed model are shown. In **A-G**) n=20 HD_S_, n=26 CI participants.

These analyses show that HD_S_ are associated with a functional skewing upon mRNA vaccination. However, functional responses quantitatively and qualitatively converge between cohorts after the third dose.

### Associations between RBD^+^ B cell and S-specific CD4^+^ T cell responses appear late in people on hemodialysis

We next examined temporal associations between B and CD4^+^ T cell responses. The net magnitudes of baseline, D1 and D2 responses for each AIM and ICS cluster were correlated with post-boost RBD^+^ B cell responses at D2 (Fig. 5A, Fig. S5A) or D3 (Fig. 5B, Fig. S5B). No positive correlation between the two cellular compartments was found at D2 in HD_S_ (Fig. 5A), in contrast to CI (Fig. S5A and (Nayrac et al., 2022)). Instead, several temporal associations were found in HD_S_ between T_H_ and RBD^+^ B cell responses at D3 (Fig. 5B). Among the subsets with significant correlations, we found clusters enriched in CXCR3 and/or CXCR5 (AIM C8, C9, C11, and C14), and functional clusters enriched in IL-2 and TNFα (ICS C4, C5, and C6) (Fig. 5B).

**Figure 5.**
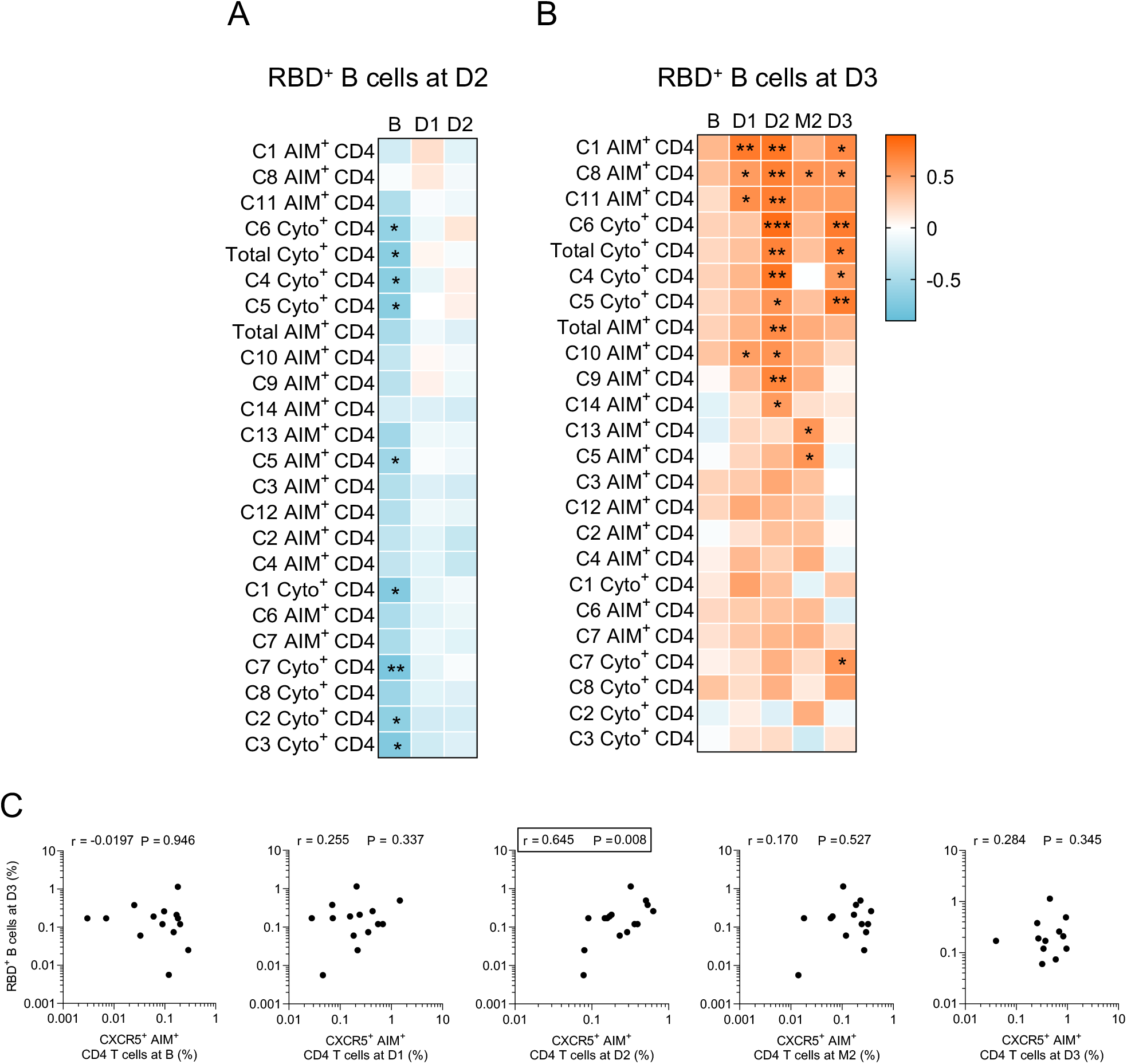
Associations between RBD^+^ B cell and S-specific CD4^+^ T cell responses appear late in people on hemodialysis. Temporal relationships between S-specific-CD4^+^ T cells and RBD^+^ B cells in HD_S_. (**A**) Correlation between total CD4^+^ T cell frequencies at B-D2 and RBD^+^ B cell frequencies at D2 in HD_S_ (n=20). **(B)** Correlation between total CD4^+^ T cell frequencies at B-D3 and RBD^+^ B cell frequencies at D3 in HD_S_ (n=20). Asterisks indicate statistically significant p values from a Spearman test (* p < 0.05, ** p < 0.01, *** p < 0.001). Colors indicate Spearman r. **(C)** Correlations between frequencies of AIM^+^ CXCR5^+^ CD4^+^ T cells (for cTfh) at the B–D3 visits and RBD^+^ B cell frequencies at D3. The r and p values from a Spearman test are indicated in each graph.

We previously reported in a low-risk population of vaccinees that S-specific CXCR5^+^ AIM^+^ CD4^+^ T cells (cT_FH_) after the first vaccine dose were predictive of RBD^+^ B cell responses after the second dose (Nayrac et al., 2022). We observed in HD_S_ an association between the cT_FH_ frequencies after the second dose and RBD^+^ B cell responses after the third dose (Fig. 5C).

These data demonstrate that like in CI, there are temporal associations between CD4^+^ T cell help and B cell responses in HD_S_ participants. However, these correlations mostly emerge after the third dose, consistent with the delayed kinetics of vaccine response in HD_S_ individuals.

### Trajectories of vaccine features highlight the need for multiple boosts in people on hemodialysis

Our data reveal multiple immune features whose trajectories differed between cohorts. To compare these trajectories, we used a normalization strategy to allow comparisons among features irrespective of their magnitude. First, we calculated the average response per participant at all timepoints, this for each feature. The ratio of the measured parameter at the timepoint to its averaged value defined its trajectory. Each ratio was then plotted on a heatmap, and clustered according to their normalized trajectory (Fig. 6AB). Three patterns were observed among HD_S_ (Fig. 6A): a first group of features peaked early after the priming (AIM^+^ C1, C12 and ICS^+^ C3). A second group showed strong responses after different boost. This group notably included humoral RBD^+^ IgG responses, CXCR3^+^ (C2, C8, C9 and C14) AIM clusters, and the TNFα^+^ IL-2^+^-enriched C6 cluster. The third pattern corresponded to late response peaking at D3 and included RBD^+^ B cells, AIM^+^ CD8^+^ T cells, total AIM^+^ and ICS^+^ CD4^+^ T cells and several AIM^+^ and ICS^+^ clusters.

**Figure 6.**
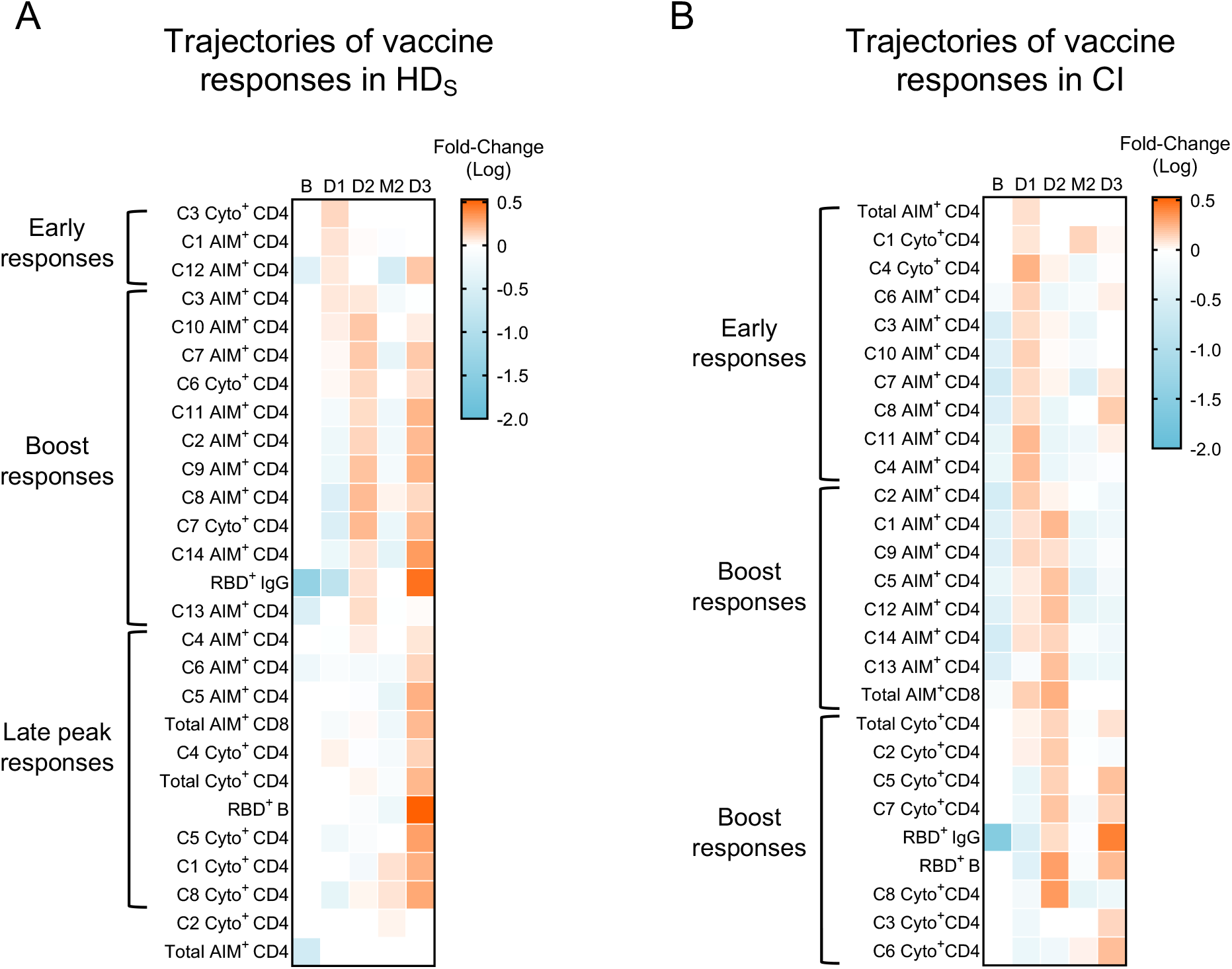
Trajectories of vaccine features highlight the need for multiple boosts in people on hemodialysis. Trajectories of specific responses to mRNA vaccines in **(A)** HD_S_ and **(B)** CI. Trajectories are represented by the fold-change of the response at each time point for a given feature, to the mean response of every time point for the same feature. Significant fold-change are colored in either orange (if increased) or blue (if decreased) (Mixed-effect analysis with Bonferroni correction for multiple comparisons, colors represent adjusted p<0.05). White color represents fold-changes that are not significantly different to the mean response. n=20 HD_S_ and 26 CI.

Different trajectories were observed in CI (Fig. 6B). Unlike HD_S_, total CD4 and CD8 responses, along with most AIM^+^ clusters, were mobilized early at D1 and were boosted at D2. The first boost enhanced total ICS^+^ CD4^+^ T, B cell and IgG responses and most ICS^+^ clusters. The second boost further recalled these responses. In contrast, these immune features were delayed in HD_S_ and mobilized only at D3. These results highlight the necessity for repeated boosting in HD_S_ to achieve peak immune responses for all immune features. This contrasts with overall earlier peak immune responses in CI, for whom the immune parameters are robustly generated after the first or the second dose.

## DISCUSSION

HD patients are at risk for severe infectious diseases, including COVID-19, and frequently poorly respond to standard vaccinations, including the initial two-dose series of SARS-CoV-2 mRNA vaccines. We show that administration of a third vaccine dose is pivotal in stimulating B cell expansion and maturation to levels similar to controls. While previous studies reported reduced IFNγ-producing T cell responses, high-dimensional functional assays demonstrate that T_H_ responses in HD are phenotypically and functionally skewed, not quantitatively inferior. Our results on cellular immunity are consistent with vaccination strategies previously proven effective in HD: administration of multiple and/or higher injections can counterbalance their low responses to immunization (Krueger et al., 2020). We show that optimal vaccine dosing interval is population-dependent: in contrast to the general population (Hall et al., 2022; Nicolas et al., 2022; Payne et al., 2021), increasing the time between the first two doses resulted in weaker humoral and cellular immunity in HD.

The third dose led to partially converging antibody levels between cohorts, although they remained lower in HD than CI at all time points. This is consistent with studies on HBV, hepatitis C virus (HCV) and influenza vaccines (Chang et al., 2012; DaRoza et al., 2003; Ghadiani et al., 2012), and previous SARS-CoV-2 studies (Grupper et al., 2021; Stumpf et al., 2021; Swai et al., 2022). The low frequencies of RBD^+^ B cell responses observed in HD after the first two doses are likely major contributors to this disparity, but their quality may play a role as well. There was delay in the maturation of B cell responses with the persistence of immature and unswitched IgM^+^ and IgD^+^ RBD^+^ B cells in HD after two mRNA vaccine doses. These features were reported in kidney transplant recipients and dialysis patients (Lederer et al., 2022; Rincon-Arevalo et al., 2021) and attributed to chronic inflammation caused by uremia toxins, along with defects of innate and T cell immunity (Betjes, 2020; Cohen, 2020; Girndt et al., 2020; Valentini et al., 2022). We also observed incomplete B cell maturation in a cohort of CI vaccinated with the standard 3-week short-interval regimen of mRNA vaccine (Nicolas et al., 2022), and thus cannot univocally delineate such defects in the 5-week interval regimen applied to the HD_S_ cohort.

Another key finding is that antigen-specific CD4^+^ T cell responses in HD were robust. Their magnitude, as measured by multiplexed AIM and ICS comparable to greater than those measured in the controls, depending on the timepoint considered. These responses were as highly diverse in phenotype and function in HD_S_ as in CI, but with qualitative differences that persisted throughout the longitudinal follow-up. We identified a pro-inflammatory/activated skewing of T_H_ responses in HD, with in CCR6, CXCR6 and HLA-DR overexpression. Such CD4^+^ T cell populations have been described as preferentially recruited at sites of inflammation in autoimmune diseases, including inflammatory kidney disease (Hou and Yuki, 2022; Linke et al., 2022). The simultaneous overexpression of the inhibitory checkpoint PD-1 by these cells may contribute to suboptimal help to other immune subsets. PD-1 upregulation on both CD4^+^ and CD8^+^ T cell populations in HD was previously reported, indicating that this dysregulation is not unique to SARS-CoV-2-specific responses (Hartzell et al., 2020).

CD4^+^ T cell responses in HD also presented functional skewing, with overrepresentation of TNFα^+^ and IL-2^+^, at the expense of IL-10^+^ and IFNγ^+^ cells. These patterns raise questions about the impact of these cytokines in the establishment of vaccine responses (Pahl et al., 2010). It has been shown that high levels of TNFα in COVID-19 could induce downstream activation of T_H1_ cells and block the final step of cT_FH_ differentiation (Kaneko et al., 2020). This skewing may contribute to the delay observed in B cell responses to SARS-CoV-2 vaccination due to an insufficient feedback inhibition of pro-inflammatory cytokines (Perianayagam et al., 2002; Ulrich et al., 2020). The third dose was characterized in HD by the normalization of the effector function profile compared to the CI. Therefore differences in the assays used (*e.g*., high-dimensional flow cytometry versus IFNγ Elispot) likely explain discrepancies between our data, in which we found robust CD4^+^ T cell responses in HD, and studies showing weaker T cell responses in this population (Broseta et al., 2021; Melin et al., 2021; Simon et al., 2022).

CD8^+^ T cell responses tend to be generated in HD only after the third dose of vaccine, in light with previous results showing that people with end-stage renal disease have more exhausted and anergic CD8^+^ T cells than CI (Hartzell et al., 2020). In both cohorts, CD8^+^ T cell responses remain low compared to their CD4^+^ T cell counterpart, consistent with the TH-biased profile of responses elicited by SARS-CoV-2 mRNA vaccines (Corbett et al., 2020; Graham, 2020).

Our results show that a long 12-week interval between the first two doses is not beneficial for people receiving hemodialysis: both B cell and antibody responses in HD_L_ after the second dose tend to be weaker to those observed in the HD_S_ cohort. The optimal dosing interval in people receiving hemodialysis remains uncertain; another study suggests that a slightly longer interval (up to 45 days, compared to 35 days in our study) was associated with stronger humoral responses (Haarhaus et al., 2022).

VOCs are an evolving challenge. After the third dose, HD developed B and CD4^+^ T cell responses specific to SARS-CoV-2 cross-reactive to Omicron, and at levels similar to CI. These findings complement previous reports showing that a third vaccine dose in HD enhanced neutralizing capacity of antibody responses against VOCs (Herman-Edelstein et al., 2022). Therefore, they might have similar protection against VOCs like Omicron than CI (Goel et al., 2022; Liu et al., 2022; Schmidt et al., 2022).

The global immune profiles observed longitudinally are consistent with a model in which HD respond more slowly to vaccination, with a third dose required to achieve B and T cell responses quantitatively and qualitatively close to those generated after two doses in CI. Temporal associations between SARS-CoV-2-specific CD4^+^ T cell and RBD^+^ B cell responses are “shifted” by one dose in HD, with a delayed link between the two features compared to CI. As cellular immune responses are comparatively less impacted by the third dose in CI than HD individuals, responses from both cohorts globally converged after the full vaccination course. Some studies have highlighted such convergence for anti-RBD IgG responses between HD and CI after three mRNA vaccine doses (El Karoui et al., 2022; Shashar et al., 2022; Verdier et al., 2022). We believe that the finding that low cellular immunity responsiveness in HD can be overcome by repeat dosing is a major positive conclusion of our study and provides an immunological basis for previous findings on the antibody responses elicited by a third dose in this vulnerable population.

Determining whether the qualitative skewing of CD4^+^ T cell responses observed in HD can alter protection against breakthrough infection, how long cellular responses persist after the third dose, and how additional booster doses can further modulate the immune profiles identified will warrant further study.

## LIMITATIONS

Our study identified several alterations of adaptive immunity elicited by SARS-CoV-2 vaccines in HD patients, with a spectrum of responsiveness in this population. Further studies are needed to better understand what individual factors may contribute to this heterogeneity. Mechanistic studies of related immune defects are very challenging, as no animal model for long-term chronic hemodialysis exists.

HD individuals are known to have frequent comorbidities and in our study were older than CI. These factors might impact immune responses independently of hemodialysis. However, while the cohorts studied are too small to conduct multivariate analyses, chronic diseases are highly prevalent in HD patients, therefore distinguishing individual factors would have limited practical impact.

This study focuses on SARS-CoV-2 naïve individuals. Additional studies are required to evaluate how prior infection shapes hybrid immunity in HD upon vaccination.

The size of the long-interval hemodialyzed cohort is small. It was not possible to recruit more suitable participants, as standard of care shifted to a short-interval regimen soon after the vaccination campaign began.

## ACKNOWLEDGMENTS

The authors are grateful to the study participants. We thank the CRCHUM BSL3 and Flow Cytometry platforms for technical assistance, Dr. Johanne Poudrier for advice and discussions. This work was supported by a FRQS Merit Research Scholar award #268471 (D.E.K), the Réseau rénal Québécois, the Fondation du CHUM, le Ministère de l’Économie et de l’Innovation du Québec, Programme de soutien aux organismes de recherche et d’innovation (A.F), a Canadian Institutes of Health Research (CIHR) operating grant # 178344 (D.E.K and A.F), a CIHR Rapid Research COVID-19 funding opportunity grant #447760 (R.S.S), a foundation grant #352417 (A.F), a CIHR operating Pandemic and Health Emergencies Research grant #177958 (A.F), and an Exceptional Fund COVID-19 from the Canada Foundation for Innovation (CFI) #41027 to A.F and D.E.K. The Symphony flow cytometer was funded by a John R. Evans Leaders Fund from the Canada Foundation for Innovation (#37521 to D.E.K) and the Fondation Sclérodermie Québec. A.F is the recipient of Canada Research Chair on Retroviral Entry no. RCHS0235 950-232424. A.P holds a Canada Research Chair in Multiple Sclerosis and the Power Corporation of Canada Chair of Université de Montréal. C.T holds the Pfizer/Université de Montréal Chair on HIV translational research. V.M.L is supported by a FRQS Junior 1 salary award. G.S is supported by a FRQS doctoral fellowship and by a scholarship from the Department of Microbiology, Infectious Disease, and Immunology of the University of Montreal. M.B was supported by a CIHR fellowship. The funders had no role in study design, data collection and analysis, decision to publish, or preparation of the manuscript.

## AUTHOR CONTRIBUTIONS

Conceptualization, G.S., A.N., M.D., A.F., R.S.S. and D.E.K.; Methodology, G.S., A.N., M.D., M.N., J.N., A.F. and D.E.K.; Software, O.T. and G.S.; Formal Analysis, J.B. and R.L.B.; Investigation, G.S., A.N., L.M., M.N., M.L., A.T., M.B., S.Y.G., C.B., G.G.L. and H.M.; Resources, M.L., R.C., A.S.F., C.B., G.G.L., H.M., N.B., G.G.O.D., L.G., C.M., P.A., C.T., V.M.L., G.G. and S.D.; Writing – Original Draft, G.S., A.N., M.D. and D.E.K.; Writing – Review & Editing, G.S., A.N., M.D. and D.E.K.; Supervision, D.E.K., A.F. and R.S.S.; Funding Acquisition, D.E.K., A.F. and R.S.S.

## STAR METHODS

### RESOURCE AVAILABILITY

#### Lead contact

Further information and requests for resources and reagents should be directed to and will be fulfilled by the lead contact, Daniel E. Kaufmann.

#### Materials availability

All unique reagents generated during this study are available from the Lead contact upon a material transfer agreement (MTA).

#### Data and code availability

The published article includes all datasets generated and analyzed for this study. Further information and requests for resources and reagents should be directed to and will be fulfilled by the Lead Contact Author.

#### Experimental Model and Subject Details

Subject characteristics are summarized in Table 1. Patients with end-stage renal disease receiving hemodialysis (HD) were enrolled into the Quebec Renal Network (QRN) COVID-19 Study as previously described (Goupil et al., 2021) and followed every 2-3 days at the *Centre Universitaire de Santé McGill* (CUSM), the *Centre Hospitalier de l’Université de Montréal* (CHUM), and the *Hôpital du Sacré-Coeur de Montréal* (HSCM). Participants from this cohort were followed and sampled before and after vaccination. Blood draws were performed at baseline (B) 12 days before first dose of vaccine with mRNA vaccine, 4 weeks after the first dose (D1), 4 weeks post second dose (D2), 12 weeks after the second dose (M2) and 4-5 weeks post third dose (D3).

Hemodialyzed participants were divided into two cohorts: a short interval cohort for which the first two vaccine doses were administered 5 weeks apart (HD_S_, n=20); and a long interval cohort (HD_L_, n=7) for which the first two doses were given 12 weeks apart, when vaccine scarcity was limiting.

The cohort of control individuals (CI, n=26) consisted of health care workers who did not have a major medical precondition qualifying for a short interval schedule (e.g, immunosuppression) and who received the first two vaccine doses 16 weeks apart per Quebec public health guidelines early in the vaccination campaign in Canada. The third inoculation was given 7 months after the second dose. Blood draws were performed at baseline (B) 1 day before the first dose of mRNA vaccine, 3 weeks after the first dose (D1), 3 weeks following the second dose (D2), 16 weeks after the second dose (M2), and 4 weeks after the third dose (D3).

Median age and interquartile range for the HD_S_ cohort was 61 [55-64], and 13 individuals were males (65%). Median age and interquartile range for the CI cohort was younger (median = 51 [41-56], p < 0.001), and 11 individuals were males (42%). The HD_L_ cohort was not significantly older (median = 66 [55-77]) and 4 individuals were males (57%). Time on hemodialysis between each cohort was comparable (See Table 1).

### METHODS DETAILS

#### Peripheral blood mononuclear cells (PBMCs) and plasma isolation

PBMCs were isolated from blood samples by Ficoll density gradient centrifugation and cryopreserved in liquid nitrogen until use. Plasma was stored at −80°C. For antibody assays, plasma was heat-inactivated for 1 hour at 56°C prior to experiments. Plasma from uninfected donors collected before the pandemic were used as negative controls and used to calculate the seropositivity threshold in our ELISA assay.

#### Enzyme-Linked Immunosorbent Assay

The SARS-CoV-2 RBD ELISA assay used was previously described (Beaudoin-Bussieres et al., 2020; Prevost et al., 2020). Briefly, recombinant SARS-CoV-2 RBD proteins or bovine serum albumin (BSA) (2.5μg/mL) as a negative control were prepared in PBS and adsorbed to plates overnight at 4°C. Coated wells were subsequently blocked with blocking buffer then washed. CR3022 mAb (50ng/ml) or a 1/250 dilution of plasma from HD, or CI donors were prepared in a diluted solution of blocking buffer and incubated with the RBD-coated wells. Plates were washed followed by incubation with the respective secondary Abs. The binding of CR3022 IgG was quantified with HRP-conjugated antibodies specific for the Fc region of human IgG and used to normalize the RLU from each plate. HRP enzyme activity was determined after the addition of a 1:1 mix of Western Lightning oxidizing and luminol reagents (Perkin Elmer Life Sciences). Light emission was measured with a LB942 TriStar luminometer (Berthold Technologies). Signal obtained with BSA was subtracted for each plasma and was then normalized to the signal obtained with CR3022 present in each plate. The seropositivity threshold was established using the following formula: mean of prepandemic SARS-CoV-2 negative plasma + (3 standard deviation of the mean of prepandemic SARS-CoV-2 negative plasma).

#### RBD-specific B cells staining

PBMCs were resuspended at 4×10^6^ cells/mL in RPMI (Gibco by Life Technologies) supplemented with penicillin/streptomycin (Gibco by Life Technologies) 10% heat inactivated FCS and incubated at 37°C, 5% CO2 for 2hrs in the presence of fluorescently labeled CCR10 antibody.

For surface stain, PBMCs were first stained for viability dye (Aquavivid, Thermofisher, 20min, 4°C) next with a mix containing a brilliant stain buffer (BD Biosciences), the surface markers for B cells detection (CD19, CD20, CD21, IgM and IgD), B cells memory phenotype (CD24, CD27, IgG and IgA), plasmablasts and plasma cells (CD38 and CD138) phenotypes, T-cells and monocytes exclusion (CD3, CD56, CD14 and CD16) (30min, 4°C) (see Table S1 for antibodies), as well as fluorescently-labeled probes for RBD^+^ B cells detection targeting two different epitopes of the RBD (RBD1-AF488 and RBD2 AF594). Omicron-RBD peptide (Accrobiosystem) was labeled, and the Omicron-RBD probe was also added into the mix where appropriate (RBD Omicron AF647). Cells were fixed with 1% paraformaldehyde (Sigma-Aldritch) for 15min at room temperature before filtration for acquisition on a FACSymphony™ A5 Cell Analyzer (BD Biosciences) and analyzed using FlowJo (BD, v10.6.2).

#### Activation induced markers (AIM) assay

PBMCs were plated in a 96-wells flat bottom plate, at 10×10^6^ cells/mL RPMI (Gibco by Life Technologies) supplemented with penicillin/streptomycin (Gibco by Life Technologies) 10% heat inactivated FCS and incubated at 37°C, 5% CO2. After a rest of 3hrs, a CD40 blocking antibody (Miltenyi) was added to the culture to prevent the interaction of CD40L with CD40 and its subsequent downregulation. In addition, antibodies for chemokine receptors CXCR6, CXCR3, CXCR5 and CCR6 were added into culture. After 15min incubation at 37°, 5% CO2, cells were stimulated with 0.5μg/mL staphylococcal enterotoxin B (SEB) or 0.5μg/mL of overlapping peptide pools for Wuhan-1 or Omicron BA.1 variants SARS-CoV-2 Spike (JPT) for 15hrs at 37°C, 5% CO2. An unstimulated condition with 0.4μL of DMSO served as a negative control.

Cells were stained for viability dye (Aquavivid, Thermofisher, 20min, 4°C), surface markers (30min, 4°C) (see Table S2 for antibodies) and fixed using 2% paraformaldehyde (Sigma-Aldritch, 15min, RT) before filtration for acquisition on the flow cytometer (FACSymphony™ A5 Cell Analyzer; BD Biosciences) and analyzed using FlowJo (BD, v10.6.2). For phenotypic analysis of antigen-specific CD4^+^ T cells, only responses that were >2-fold over unstimulated condition were included to limit the impact of background staining. In contrast, for analysis of antigen-specific CD4^+^ T cells subsets as percentage of total CD4^+^ T cells, background-subtracted net values were used, which did not require excluding responses.

#### Intracellular cytokines staining (ICS) assay

PBMCs were resuspended at 10×10^6^ cells/mL RPMI (Gibco by Life Technologies) supplemented with penicillin/streptomycin (Gibco by Life Technologies) 10% heat inactivated FCS and incubated at 37°C, 5% CO2. After a rest of 2hrs, cells were stimulated with 0.5μg/mL staphylococcal enterotoxin B (SEB) or 0.5μg/mL of overlapping peptide pools for Wuhan-1 or Omicron BA.1 variants SARS-CoV-2 Spike (JPT) for 6hrs at 37°C, 5% CO2. An unstimulated condition with 0.4μL of DMSO served as a negative control. Brefeldin A (BD Biosciences), Monensin-1 (BD Biosciences) and a fluorescently labeled CD107a antibody were added for the remaining 5hrs.

Cells were stained for viability dye (Aquavivid, Thermofisher, 20min, 4°C), surface markers (30min, 4°C) and intracellularly for cytokines (30min, room temperature) using the IC Fixation/Permeabilization kit (eBioscience) (see Table S3 for antibodies) and filtrated before acquisition on the flow cytometer (FACSymphony™ A5 Cell Analyzer, BD Biosciences) and analyzed using FlowJo (BD, v10.6.2).

#### Statistics

Symbols represent biologically independent samples from HD and CI donors. Lines connect data from the same donor. Thick lines represent median values. Linear mixed models fitting cell frequencies in terms of cohort, time point and their interaction were run using R and the package “nlme”. Model diagnostics were performed, checking for heteroscedasticity and normality among residuals. All retained models used a squareroot transform on the response variable, which helped in reducing the impact of outliers. Post-hoc contrasts across all pairwise comparisons of factor levels were obtained with the package “emmeans”, correcting the p values by the method of Holm-Bonferroni where applicable. An important caveat of the square-root transform is that the reported contrast estimates and their confidence intervals remain on this scale, making their interpretation tricky. This was not deemed too great an obstacle, as qualitative statements on significant contrasts could be made based on p-values. Thirty-five linear mixed models were retained, those being anti-RBD IgG, RBD B, AIM CD4, ICS CD4, AIM CD8, CXCR3, CXCR5, CXCR6, CCR6, PD-1, CD38, HLA-DR, IFNγ, IL-2, TNFα, IL-10, CD107a and IL-17A being compared between HD_S_, CI and HD_L_ cohorts. There were also comparisons of HD_S_, HD_L_ and CI for anti-RBD IgG, RBD B, AIM CD4, ICS CD4 and AIM CD8. Models without satisfactory diagnostics were abandoned in favor of non-parametric methods. Differences in responses for the same patient before and after vaccination were performed using Wilcoxon matched pair tests. Differences in responses between HD_S_ and CI were measured by Mann-Whitney tests. Wilcoxon and Mann-Whitney tests were generated using GraphPad Prism (version 9.2.0). p values <0.05 were considered significant. p values are indicated for each comparison assessed. For descriptive correlations, Spearman’s R correlation coefficient was applied. For graphical representation on a log scale (but not for statistical tests), null values were arbitrarily set at the minimal values for each assay.

#### Software scripts and visualization

Graphics and pie charts were generated using GraphPad PRISM (v9.2.0) and ggplot2 (v3.3.3) in R (v4.1.0). Heat maps were generated in R (v4.1.0) using the *pheatmap* package (v1.0.12). Principal component analyses were performed with the *prcomp* function (R). Uniform manifold approximation and projection (UMAP) was performed using package M3C (v1.14.0) on gated FCS files loaded through the flowCore package (v2.4.0). Samples were downsampled to a comparable number of events (300 cells for AIM, 100 cells for ICS). Scaling and logical transformation of the flow cytometry data was applied using the FlowSOM (Quintelier et al., 2021) R package (v2.0.0). All samples at all time points were loaded. Clustering was achieved using Phenograph (v0.99.1) with the hyperparameter k (number of nearest neighbors) set to 150). We previously provided all R codes scripted for this paper in another study (Nayrac et al., 2022). We obtained an initial 14 AIM^+^ and 8 ICS^+^ clusters. For B and CD4^+^ T cell phenotyping, only participants with ≥5 RBD^+^ B events across all depicted time points were analyzed.

