## Supplemental Figures and Tables for "A third SARS-CoV-2 mRNA vaccine dose in people receiving hemodialysis overcomes B cell defects but elicits a skewed CD4^+^ T cell profile"

Figure S1

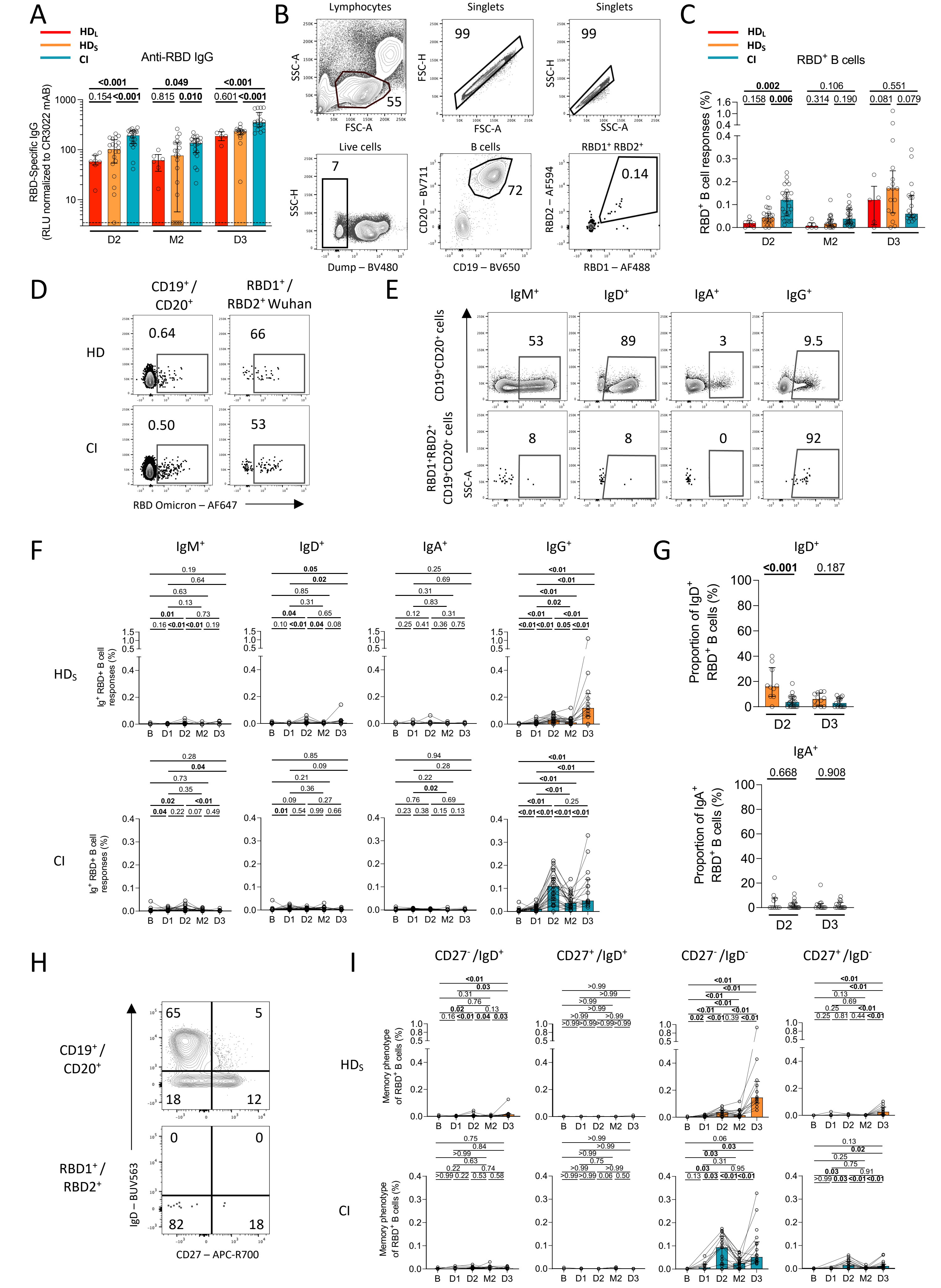

A
HD_L
HD_S
CI
Anti-RBD IgG
RBD-Specific IgG (RLU normalized to CR3022 mAB)
D2
M2
D3
B
Lymphocytes
Singlets
SSC-A
FSC-A
FSC-H
SSC-H
Live cells
B cells
RBD1⁺ RBD2⁺
Dump – BV480
CD19 – BV650
CD20 – BV711
RBD2 – AF594
RBD1 – AF488
C
RBD⁺ B cells
RBD⁺ B cell responses (%)
D
CD19⁺ / CD20⁺
RBD1⁺ / RBD2⁺ Wuhan
HD
CI
RBD Omicron – AF647
E
IgM⁺
IgD⁺
IgA⁺
IgG⁺
CD19⁺CD20⁺ cells
RBD1⁺RBD2⁺ CD19⁺CD20⁺ cells
SSC-A
F
HDs
CI
Ig⁺ RBD+ B cell responses (%)
B
D1
D2
M2
D3
G
IgD⁺
IgA⁺
Proportion of IgD⁺ RBD⁺ B cells (%)
Proportion of IgA⁺ RBD⁺ B cells (%)
H
CD19⁺ / CD20⁺
RBD1⁺ / RBD2⁺
IgD – BUV563
CD27 – APC-R700
I
CD27⁻ /IgD⁺
CD27⁺ /IgD⁺
CD27⁻ /IgD⁻
CD27⁺ /IgD⁻
Memory phenotype of RBD⁺ B cells (%)

Figure S2

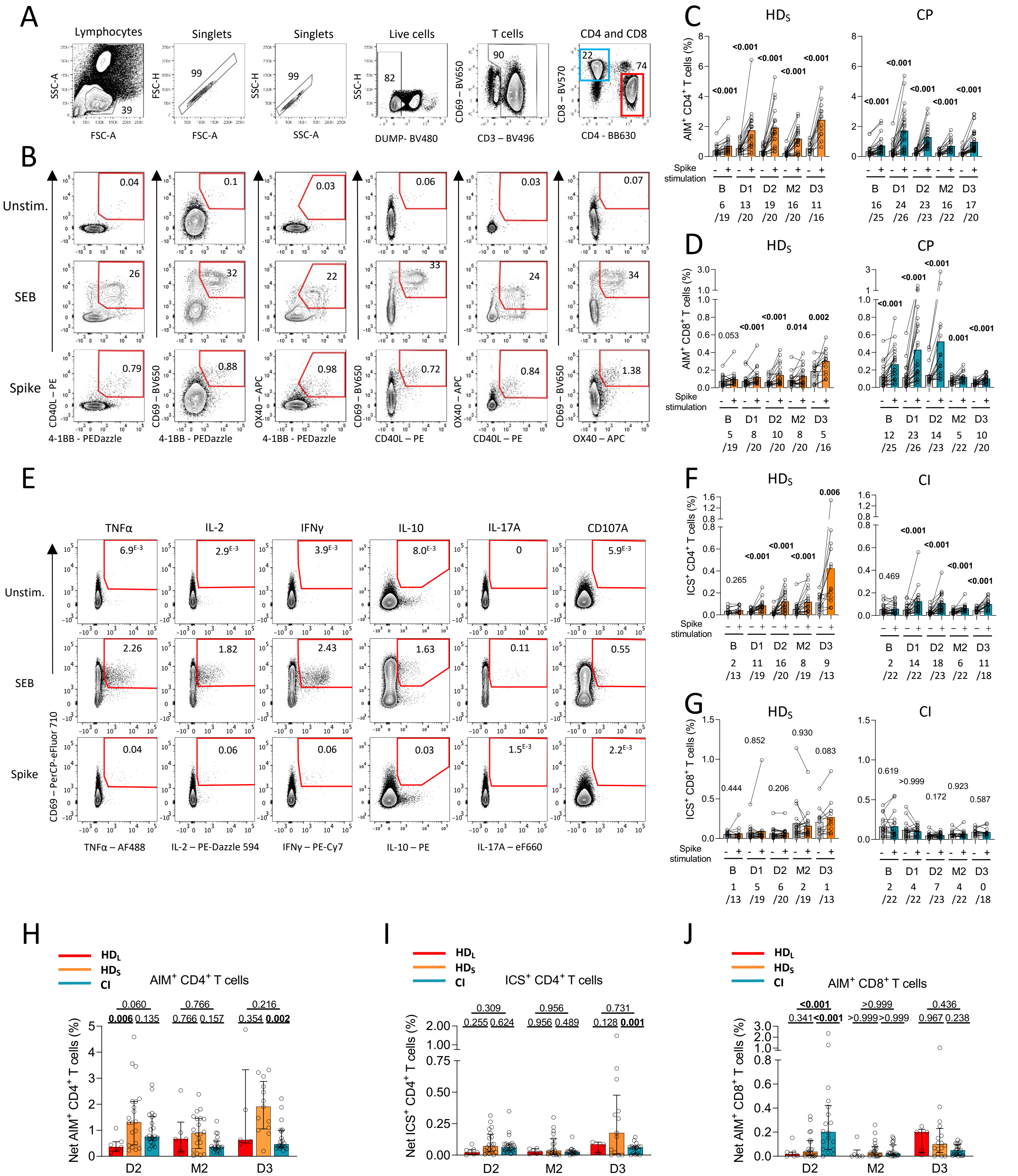

A
Lymphocytes
Singlets
Singlets
Live cells
T cells
CD4 and CD8
SSC-A
FSC-A
FSC-H
SSC-H
DUMP- BV480
CD69 – BV650
CD3 – BV496
CD8 – BV570
CD4 - BB630
39
99
99
82
90
22
74
B
Unstim.
SEB
Spike
0.04
0.1
0.03
0.06
0.03
0.07
26
32
22
33
24
34
0.79
0.88
0.98
0.72
0.84
1.38
CD40L – PE
CD69 – BV650
OX40 – APC
CD69 – BV650
OX40 – APC
CD69 – BV650
4-1BB - PEDazzle
4-1BB - PEDazzle
4-1BB - PEDazzle
CD40L – PE
CD40L – PE
OX40 – APC
C
HDs
CP
AIM+ CD4+ T cells (%)
Spike stimulation
B D1 D2 M2 D3
6 /19 13 /20 19 /20 16 /20 11 /16
16 /25 24 /26 23 /23 16 /22 17 /20
D
HDs
CP
AIM+ CD8+ T cells (%)
Spike stimulation
B D1 D2 M2 D3
5 /19 8 /20 10 /20 8 /20 5 /16
12 /25 23 /26 14 /23 5 /22 10 /20
E
TNFα
IL-2
IFNγ
IL-10
IL-17A
CD107A
Unstim.
SEB
Spike
CD69 – PerCP-eFluor 710
TNFα – AF488
IL-2 – PE-Dazzle 594
IFNγ – PE-Cy7
IL-10 – PE
IL-17A – eF660
F
HDs
CI
ICS+ CD4+ T cells (%)
Spike stimulation
B D1 D2 M2 D3
2 /13 11 /19 16 /20 8 /19 9 /13
2 /22 14 /22 18 /23 6 /22 11 /18
G
HDs
CI
ICS+ CD8+ T cells (%)
Spike stimulation
B D1 D2 M2 D3
1 /13 5 /19 6 /20 2 /19 1 /13
2 /22 4 /22 7 /23 4 /22 0 /18
H
HDL
HDS
CI
AIM+ CD4+ T cells
Net AIM+ CD4+ T cells (%)
D2 M2 D3
I
HDL
HDS
CI
ICS+ CD4+ T cells
Net ICS+ CD4+ T cells (%)
D2 M2 D3
J
HDL
HDS
CI
AIM+ CD8+ T cells
Net AIM+ CD8+ T cells (%)
D2 M2 D3

Figure S3

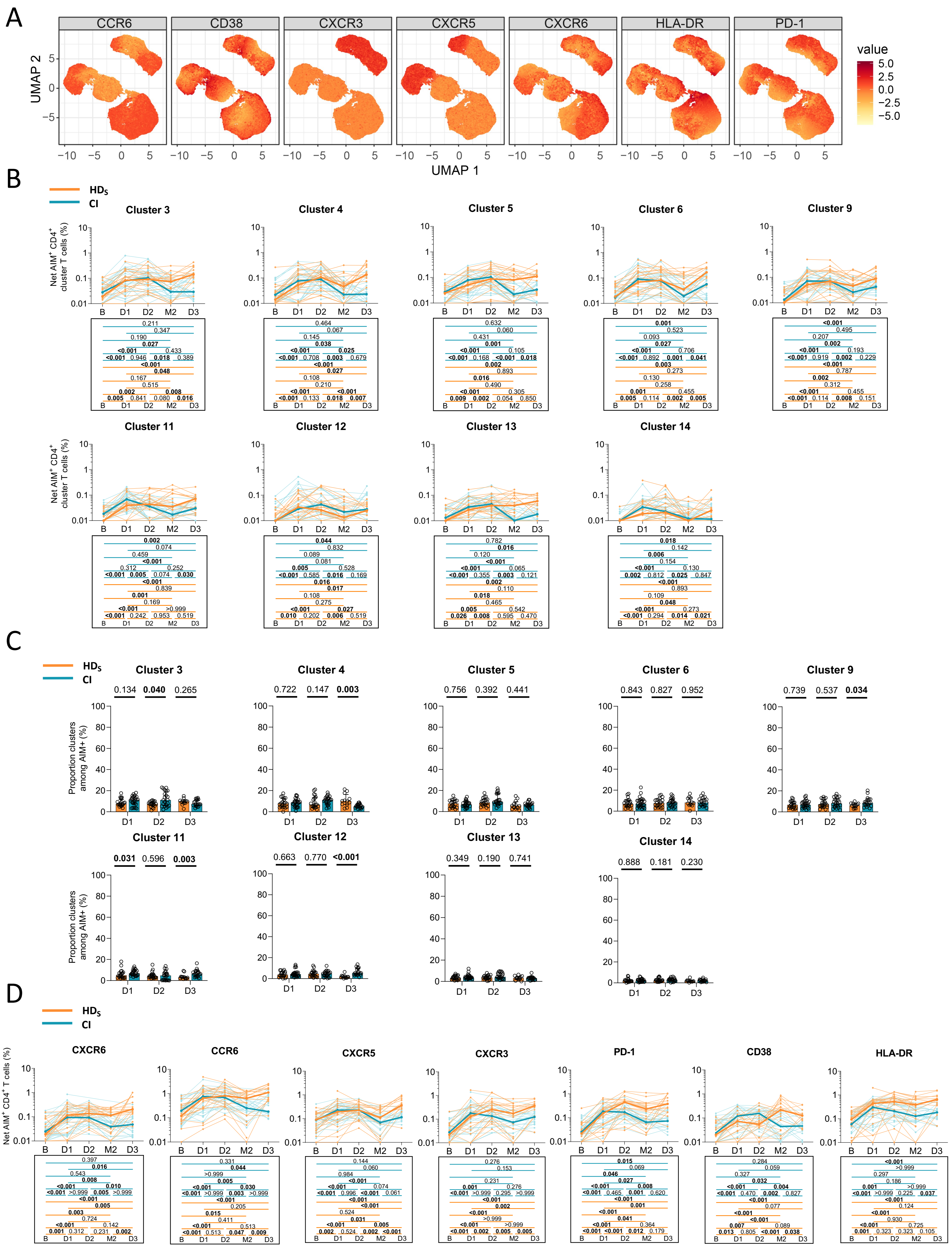

Figure S4

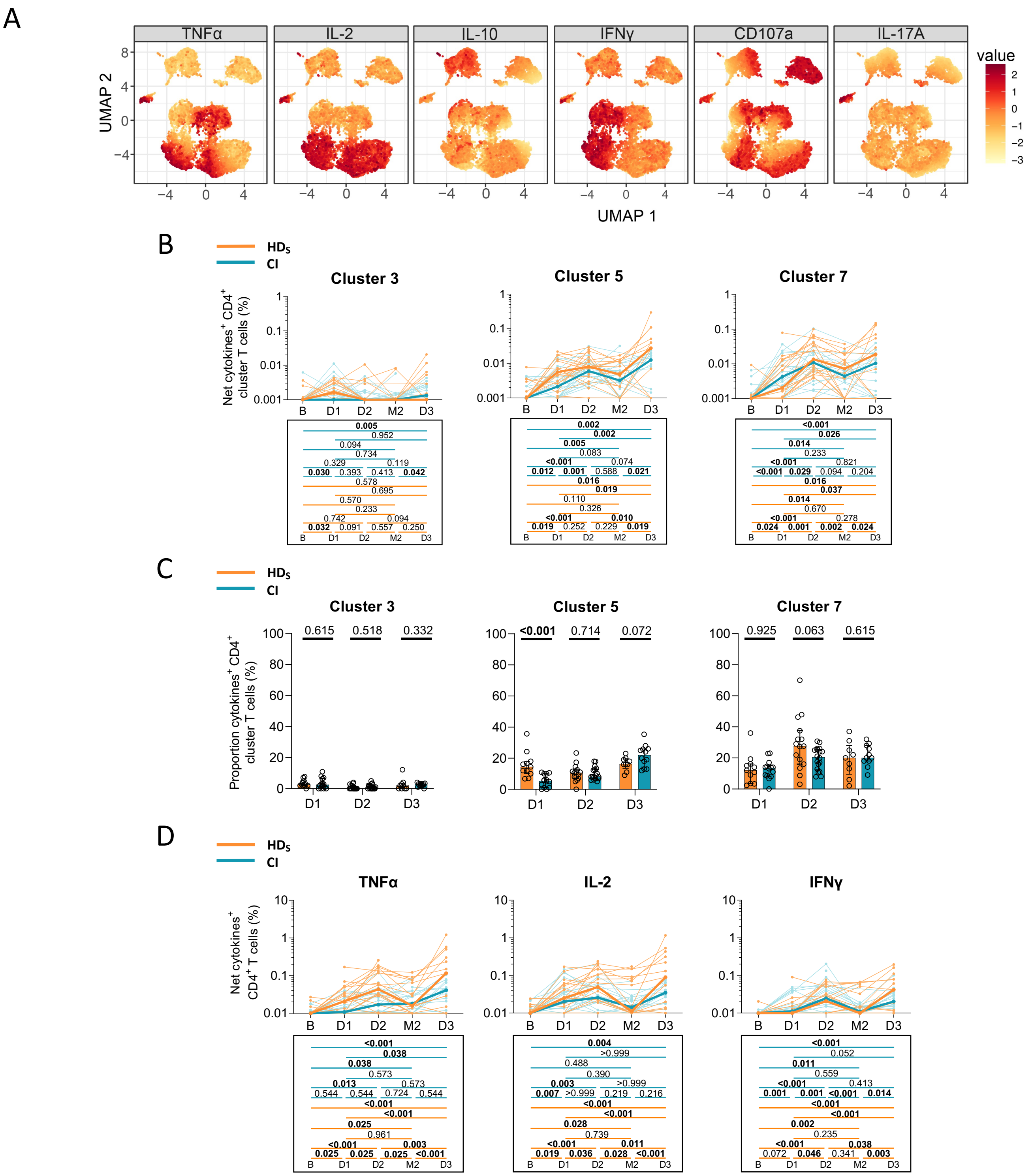
A
TNFα
IL-2
IL-10
IFNγ
CD107a
IL-17A
UMAP 2
UMAP 1
value
B
HDs
CI
Cluster 3
Cluster 5
Cluster 7
Net cytokines+ CD4+ cluster T cells (%)
C
Proportion cytokines+ CD4+ cluster T cells (%)
D
Net cytokines+ CD4+ T cells (%)
B
D1
D2
M2
D3

Figure S5

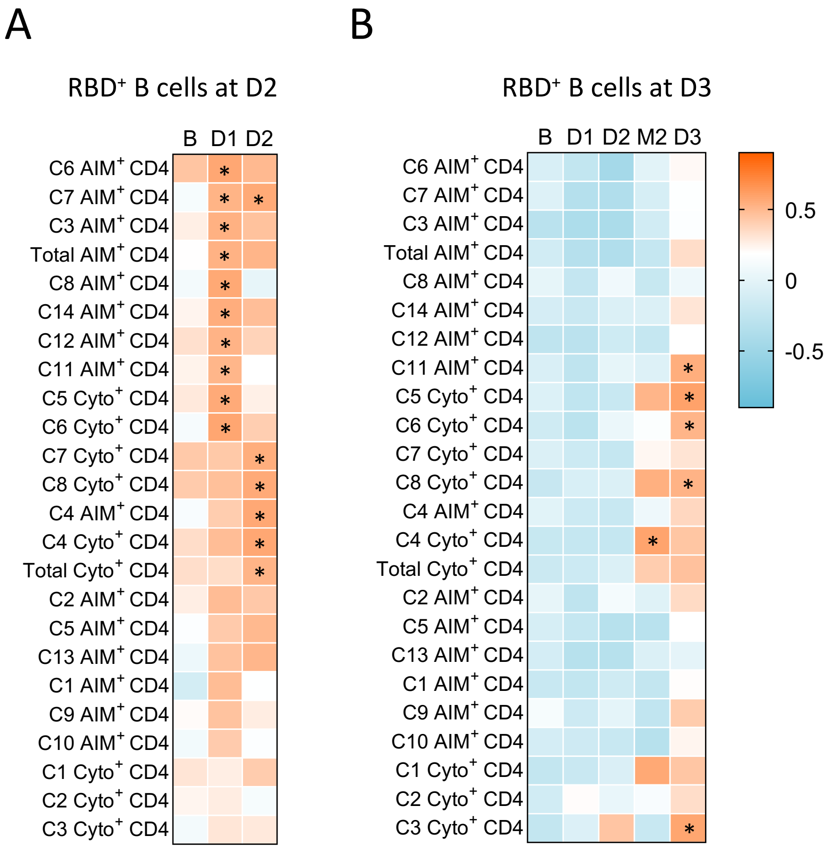
A
RBD+ B cells at D2
B D1 D2
C6 AIM+ CD4
C7 AIM+ CD4
C3 AIM+ CD4
Total AIM+ CD4
C8 AIM+ CD4
C14 AIM+ CD4
C12 AIM+ CD4
C11 AIM+ CD4
C5 Cyto+ CD4
C6 Cyto+ CD4
C7 Cyto+ CD4
C8 Cyto+ CD4
C4 AIM+ CD4
C4 Cyto+ CD4
Total Cyto+ CD4
C2 AIM+ CD4
C5 AIM+ CD4
C13 AIM+ CD4
C1 AIM+ CD4
C9 AIM+ CD4
C10 AIM+ CD4
C1 Cyto+ CD4
C2 Cyto+ CD4
C3 Cyto+ CD4
B
RBD+ B cells at D3
B D1 D2 M2 D3
C6 AIM+ CD4
C7 AIM+ CD4
C3 AIM+ CD4
Total AIM+ CD4
C8 AIM+ CD4
C14 AIM+ CD4
C12 AIM+ CD4
C11 AIM+ CD4
C5 Cyto+ CD4
C6 Cyto+ CD4
C7 Cyto+ CD4
C8 Cyto+ CD4
C4 AIM+ CD4
C4 Cyto+ CD4
Total Cyto+ CD4
C2 AIM+ CD4
C5 AIM+ CD4
C13 AIM+ CD4
C1 AIM+ CD4
C9 AIM+ CD4
C10 AIM+ CD4
C1 Cyto+ CD4
C2 Cyto+ CD4
C3 Cyto+ CD4
0.5
0
-0.5

### SUPPLEMENTARY FIGURE LEGENDS

**Figure S1. Related to Figure 1. (A)** Comparison of RBD-specific IgG responses between $HD_L$ (red), $HD_S$ (orange) and CI (blue) participants at D2, M2 and D3. Bars represent median ± interquartile range. Intercohort statistical comparisons using a linear mixed model are shown. **(B)** Gating strategy to identify $RBD^+$ B cells. **(C)** Comparison of $RBD^+$ B cell responses between $HD_L$ (red), $HD_S$ (orange) and CI (blue) participants at D2, M2 and D3. Bars represent median ± interquartile range. Intercohort statistical comparisons using a linear mixed model are shown. **(B)** Gating strategy to identify $RBD^+$ B cells. **(D)** Gating strategy of Omicron $RBD^+$ B cells among total B cells (left) and among Wuhan-1-$RBD^+$ B cells (right). **(E)** Examples of gatings for IgD, IgM, IgA and IgG expression on total $CD19^+CD20^+$ B cells (top) or $RBD^+$ B cells (bottom). **(F)** Histograms reporting the longitudinal frequency of isotype expression in HDs (orange) and CI (blue) participants. Lines connect data points for individual participants. Wilcoxon tests are shown above each panel. **(G)** Comparison of $IgD^+$ and $IgA^+$ $RBD^+$ B cells between $HD_S$ and CI participants at D2 and D3. Mann-Whitney tests are shown. **(H)** Gating strategy of $IgD^{+/-}$ and $CD27^{+/-}$ of total (left) and $RBD^+$ memory B cells (right). **(I)** Histograms reporting the longitudinal frequency of each IgD and CD27 RBD-B phenotypes in $CD19^+$ $CD20^+$ B cells for HDs (orange) and CI (blue) participants. In support to the pie charts displayed in Figure 1K. Wilcoxon tests are shown above.

**Figure S2. Related to Figure 2. (A)** Representative upstream generic gating and **(B)** ORgate strategy to identify SARS-CoV-2-specific $AIM^+$ T cells. For simplicity, the example focuses on $CD4^+$ T cells. **(CD)** Raw frequencies of **(C)** $AIM^+$ CD4+ and **(D)** $CD8^+$ T cells following *ex vivo* stimulation of PBMCs with a pool of SARS-CoV-2 Spike peptides (colored). $HD_S$ are represented on the left and CI on the right for each panel. As a control, PBMCs cells were left unstimulated (grey bars). The bars represent median values. Wilcoxon tests are shown. Number of responders that were at least two times over unstimulated conditions are written below the histograms for each timepoints. **(E)** Representative ORgate strategy to identify SARS-CoV-2-specific cytokine-expressing T cells. For simplicity, the example focuses on CD4 T cells. **(FG)** Raw frequencies of **(F)** cytokine-expressing $CD4^+$ T cells and **(G)** $CD8^+$ T cell following *ex vivo* stimulation of PBMCs with a pool of SARS-CoV-2 Spike peptides (colored). $HD_S$ are represented on the left and CI on the right for each panel. As a control, PBMCs cells were left unstimulated (grey bars). The bars represent median values. Wilcoxon tests are shown. Number of responders that were at least two times over unstimulated conditions are written below the histograms for each timepoints. **(HIJ)** Comparison at D2, M2 and D3 of **(H)** net $AIM^+$ $CD4^+$ T cell responses, **(I)** net $ICS^+$ $CD4^+$ responses and **(J)** net $AIM^+$ $CD8^+$ responses between $HD_L$ (red), $HD_S$ (orange) and CI (blue) participants. The bars represent median and interquartile ranges. Intercohort statistical comparisons using a linear mixed model. In A-G) n=20 $HD_S$, n=26 CI participants, in H-J) n=20 $HD_S$, n=26 CI, n=7 $HD_L$ participants.

**Figure S3. Related to Figure 3. (A)** Heat map overlaid on the $AIM^+$ UMAP showing the gradient of expression for each marker. **(B)** Longitudinal analysis of net $AIM^+$ $CD4^+$ T cell clusters, regarding clusters 3, 4, 5, 6, 9, 11, 12, 13, and 14 for $HD_S$ (orange, n=20) and CI (blue; n=26) participants. Lines connect data from the same donor. Bold lines represent median values. Wilcoxon tests are shown below for each pairwise comparison. Complement Figure 3E. **(C)** Proportions of $AIM^+$ clusters 3, 4, 5, 6, 9, 11, 12, 13 and 14 among $AIM^+$ $CD4^+$ T cells in $HD_S$ and CI at D1, D2 and D3. Bars represent median ± interquartile range. Mann-Whitey tests are shown. Complement Figure 3F. **(D)** Longitudinal frequencies of $CCR6^+$, $CXCR5^+$, $CXCR3^+$, PD-$1^+$, $CD38^+$ and HLA-$DR^+$ $AIM^+$ $CD4^+$ T cells in $HD_S$ (orange) and CI (blue). Lines connect data from the same donor. The bold lines represent the median value of each cohort. Statistical comparisons using a linear mixed model.

**Figure S4. Related to Figure 4. (A)** Heat map overlaid on the cytokine$^+$ UMAP showing the gradient of expression for each marker. **(B)** Longitudinal analysis of net ICS$^+$ CD4$^+$ T cell clusters, regarding clusters 3, 5 and 7 for $HD_S$ (orange, n=20) and CI (blue; n=26) participants. Lines connect data from the same donor. Bold lines represent median values. Wilcoxon tests are shown below for each pairwise comparison. Complement Figure 4E. **(C)** Proportions of ICS$^+$ clusters 3, 5 and 7 among ICS$^+$ CD4$^+$ T cells in $HD_S$ and CI at D1, D2 and D3. Bars represent median ± interquartile range. Mann-Whitey tests are shown. Complement Figure 4F. **(D)** Longitudinal frequencies of TNFα$^+$, IL-2$^+$ and IFNγ$^+$ CD4$^+$ T cells in $HD_S$ (orange) and CI (blue). Lines connect data from the same donor. The bold line represents the median value of each cohort. Statistical comparisons using a linear mixed model.

**Figure S5: Related to Figure 5.** Temporal relationships between S-specific CD4$^+$ T cells and RBD$^+$ B cells in CI. **(A)** Correlation between total CD4$^+$ T cell frequencies at B-D2 and RBD$^+$ B cell frequencies at D2 in CI (n=26). **(B)** Correlation between total CD4$^+$ T cell frequencies at B-D3 and RBD$^+$ B cell frequencies at D3 in CI (n = 26). Asterisks indicate statistically significant p value from a Spearman test ($p<0.05$). Colors indicate Spearman r. **(C)** Correlations between frequencies of AIM$^+$ CXCR5$^+$ CD4$^+$ T cells (for cTfh) at the B–D3 visits and RBD$^+$ B cell frequencies at D3. The r and p values from a Spearman test are indicated in each graph.

**Table S1. Flow cytometry antibody staining panel for B cells characterization**

| Marker-Fluorophore | Clone | Source | Catalog # |
|---|---|---|---|
| CD3 – BV480 | UCHT1 | BD Biosciences | 566105 |
| CD14 – BV480 | M5E2 | BD Biosciences | 746304 |
| CD16 – BV480 | 3G8 | BD Biosciences | 566108 |
| CD19 – BV650 | SJ25C1 | Biolegend | 363026 |
| CD20 – BV711 | 2H7 | Biolegend | 563126 |
| CD21 – BV786 | B-LY4 | BD Biosciences | 740969 |
| CD24 – BUV805 | ML5 | BD Biosciences | 742010 |
| CD27 – APC-R700 | M-T271 | BD Biosciences | 565116 |
| CD38 – BB790 | HIT2 | BD Biosciences | CUSTOM |
| CD56 – BV480 | NCAM16.2 | BD Biosciences | 566124 |
| CD138 – BUV661 | MI15 | BD Biosciences | 5 749873 |
| CCR10 – BUV395 | 1B5 | BD Biosciences | 565322 |
| HLA-DR – BB700 | G46-6 | BD Biosciences | 566480 |
| IgA - PE | IS11-8E10 | Miltenyi | 130-113-476 |
| IgD – BUV563 | IA6-2 | BD Biosciences | 741394 |
| IgG – BV421 | G18-147 | BD Biosciences | 562581 |
| IgM – BUV737 | UCH-B1 | Thermo Fisher Scientific | 748928 |
| LIVE/DEAD Fixable dead cell | N/A | Thermo Fisher Scientific | L34960 |

**Table S2. Flow cytometry antibody staining panel for activation-induced marker assay**

| Marker-Fluorophore | Clone | Source | Catalog # |
|---|---|---|---|
| CD3 – BUV496 | UCHT1 | BD | 612941 |
| CD4 – BB630 | SK3 | BD | 624294 |
| CD8 – BV570 | RPA-T8 | Biolegend | 301037 |
| CD14 – BV480 | M5E2 | BD | 746304 |
| CD19 – BV480 | HIB19 | BD | 746457 |
| CD38 – BB790 | HIT2 | BD | CUSTOM |
| CD45RA – PerCP Cy5.5 | HI100 | BD | 563429 |
| CD69 – BV650 | FN50 | Biolegend | 310934 |
| CD134 (OX40) - APC | ACT35 | BD | 563473 |
| CD137 (4-1BB) – PE-Dazzle 594 | 4B4-1 | Biolegend | 309826 |
| CD154 (CD40L) - PE | TRAP1 | BD | 555700 |
| CD183 (CXCR3) – BV605 | G025H7 | Biolegend | 353728 |
| CD185 (CXCR5) – BV421 | J25D4 | Biolegend | 356920 |
| CD186 (CXCR6) – BUV805 | 13B 1E5 | BD | 748448 |
| CD196 (CCR6) – BUV737 | 11A9 | BD | 564377 |
| CD279 (PD1) – BV711 | EH122H | Biolegend | 329928 |
| HLA-DR - FITC | LN3 | Biolegend | 327005 |
| LIVE/DEAD Fixable dead cell | N/A | Thermo Fisher Scientific | L34960 |

**Table S3. Flow cytometry antibody staining panel for intracellular cytokines staining assay**

| Marker-Fluorophore | Clone | Source | Catalog # |
|---|---|---|---|
| CD3 – BUV395 | UCHT1 | BD Biosciences | 563546 |
| CD4 – BV711 | L200 | BD Biosciences | 563913 |
| CD8 – BV570 | RPA-T8 | Biolegend | 301037 |
| CD14 – BUV805 | M5E2 | BD Biosciences | 612902 |
| CD16 – BV650 | 3G8 | Biolegend | 302042 |
| CD19 – APC-eFluor780 | HIB19 | Thermo Fisher Scientific | 47-0199 |
| CD56 – BUV737 | NCAM16.2 | BD Biosciences | 564448 |
| CD69 – PerCP-eFluor710 | FN50 | Thermo Fisher Scientific | 46-0699-42 |
| CD107A – BV786 | H4A3 | BD Biosciences | 563869 |
| IFN-γ – PECy7 | B27 | BD Biosciences | 557643 |
| CD154 (CD40L) – BV421 | TRQP1 | BD Biosciences | 563886 |
| IL-2 – PE-Dazzle 594 | MQ1-17H12 | Biolegend | 500344 |
| IL-10 - PE | JES3-9D7 | BD Biosciences | 554498 |
| IL-17A – eFluor660 | eBio64CAP17 | Thermo Fisher Scientific | 50-7179-42 |
| TNF-α – Alexa Fluor 488 | Mab11 | Thermo Fisher Scientific | 502915 |
| Granzym B – Alexa Fluor 700 | GB11 | BD | 561016 |
| LIVE/DEAD Fixable dead cell | N/A | Thermo Fisher Scientific | L34960 |
